# Bioorthogonal labeling of chitin in pathogenic Candida species reveals biochemical mechanisms of hyphal growth and homeostasis

**DOI:** 10.1101/2024.08.27.609898

**Authors:** Caroline Williams, Bella R. Carnahan, Stephen N. Hyland, Catherine L. Grimes

## Abstract

Pathogenic fungi rely on the cell wall component, chitin, for critical structural and immunological functions. Here a chitin labeling method to visualize the hyphal pathogenic response was developed. The data show that filamentous fungi, *Candida albicans*, transport *N*-acetylglucosamine (NAG) bio-orthogonal probes and incorporate them into the cell wall, indicating the probes utility for *in vivo* study of the morphological, pathogenic switch. As yeast reside in complex microenvironments, The data show that the opportunistic microbe *C. albicans*, has developed processes to utilize surrounding bacterial cell wall fragments to initiate the morphogenic switch. The probes are utilized for visualization of growth patterns of pathogenic fungi, providing insights into novel mechanisms for the development of antifungals. Remodeling chitin in fungi using NAG derivatives will advance yeast pathogenic studies.

## Main Text

Chitin is the second most abundant polysaccharide in nature after cellulose and serves as a vital biopolymer used by forms of life ranging from pathogenic fungi to crustaceans to insects (*1*). In fungi, chitin is composed of a repeating chain of β-1,4-*N*-acetyl-glucosamine (GlcNAc/NAG) linked sugars that fold onto each other to form anti-parallel chains (*2, 3*). This polymer is essential to the plasticity of the fungal cell wall and provides cells with protection from external stressors (*4*). In addition, this biopolymer, absent from mammalian cells, serves as a potent immunomodulator, activating the innate immune system by engaging with Toll-like receptors and other inflammasomes (*5*).

In the pathogenic fungi *Candida albicans* (*C. albicans*), chitin is biosynthesized de novo from glucose via a conserved linear pathway (Fig. 1) (*6, 7*). However, it can also be synthesized by sequestering exogenous GlcNAc and converting it to 6-phospho-GlcNAc via the enzyme Hexokinase-1 (Hxk1) (*6, 8*). This marks the carbohydrate for downstream use in the chitin biosynthesis pathway. The phosphoacetylglucosamine mutase, Pcm1, rearranges the phosphate group to the anomeric, 1-hydroxy position (*9*) and then Uap1, a UDP-*N*-acetylglucosamine pyrophosphorylase, converts this molecule to UDP-GlcNAc (*10*). This nucleotide sugar is a precursor for multiple molecules, but it can also be then polymerized by a series of chitin synthases (*6, 11*). In *C. albicans* there are four chitin synthases (Chs1, Chs3, Chs2, and Chs8) that work to deposit chitin at growth sites in the cell (*11*). A complementary set of enzymes - chitin degraders or chitinases – are capable of breaking down the polymer to permit rearrangement and cell growth (*1, 12*).

**Fig. 1.**
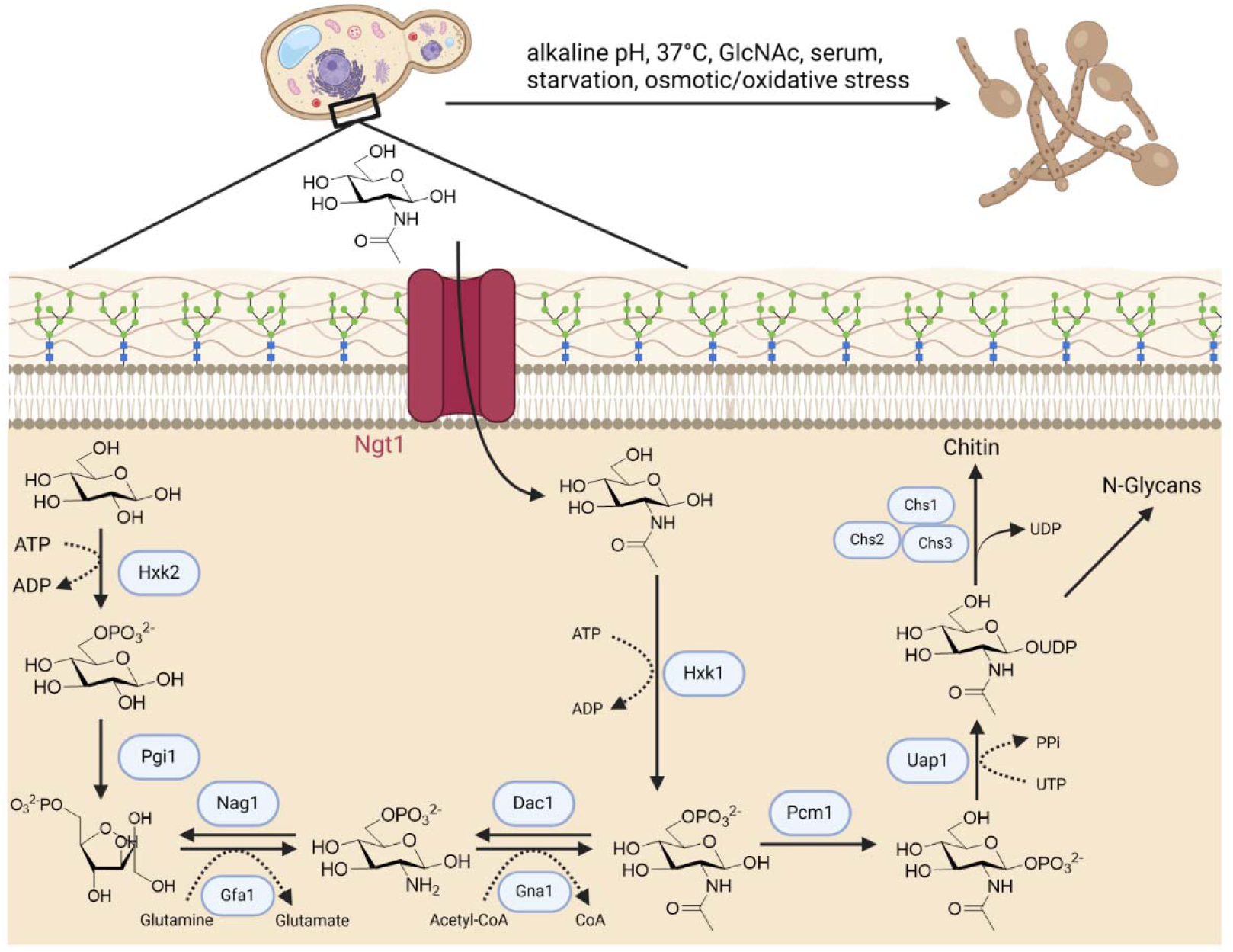
Chitin biosynthesis in *C. albicans*. Glucose or exogenous GlcNAc can be salvaged by the cell and used as precursors to chitin in fungal cells, via the GlcNAc specific transporter Ngt1 and the specific kinase Hxk1. The intermediate UDP-GlcNAc can either be shuttled to chitin or surface glycans, such as *N*-glycans.

In order to study these essential biochemical processes, a number of biochemical tools have been developed. One of the most popular methods for labeling chitin is the use of Calcofluor white, an ionic compound that coordinates with the hydroxyl groups in chitin’s backbone. It permits for visibility of the polymer using confocal microscopy; however, its mechanism of binding by competing for hydrogen binding sites, further inhibits the growth of chitin (*13*). For chitin degraders, methods involving tracking radiolabeled ^14^C substrates and colorimetric incubation experiments have been developed (*12*).

Currently, there exists no robust mechanism that labels the components involved in the remodeling of the fungal cell wall. Fungal cell wall biosynthesis and its molecular components have been thoroughly understood and characterized since the late 1970’s (*14, 15*). Motivation to utilize the biosynthetic machinery to tag the fungal cell wall is driven by prior successes in the field with in vivo labeling of macromolecular structures and polymers such as peptidoglycan (PG) (*16, 17*). Pioneering work conducted by Bertozzi introduced bio-orthogonal functionalities into eukaryotic glycans, in which a metabolic engineering strategy to label the cell wall, specifically N-glycans on the surface of the yeast cell wall was utilized (*18*). Here, the dedicated kinase, Hxk1, from *C. albicans* was encoded into the genetically tractable test strain, *Saccharomyces cerevisiae* (*S. cerevisiae*). This modified *S. cerevisiae* strain was then supplemented GlcNAc building blocks which contained a bio-orthogonal azide or alkyne handle. In this innovative strategy, Bertozzi and co-workers showed that these bio-orthogonal probes were capable of being incorporated into the fungal cell wall. However, as this was a modified strategy, the labeling was restricted to bud scars and the azide/alkyne supplemented cells showed morphological discrepancies compared to the GlcNAc fed cultures (*18*). Here, we wished to build on this method by working directly with the human pathogen, *C. albicans* and labeling chitin with modified GlcNAc carbohydrates.

*C. albicans*, a commensal fungal organism, resides on human barrier surfaces such as the skin as well as colonizes areas such as the mouth, lungs, gut, and vagina (*19-21*). During periods of dysbiosis, *C. albicans* transitions into a pathogenic state indicated by discrete morphological phases, such as budding, pseudohyphal, and hyphal forms (*22-24*). During this morphological switch, *C. albicans* upregulates its production of chitin, which normally sits at 2-3% dry weight (*25*), to sustain the increased content required in elongated hyphae (*5, 26*). This morphological flexibility is also a virulence determinant, as the switch from budding to hyphae plays a key role in the infection process by allowing penetration of the local tissues and avoidance of phagocytic destruction (*27, 28*). In order to combat infection, the human innate immune system uses a variety of pathogen recognition receptors (PRR’s) to recognize certain fingerprints (pathogen associated molecular patterns – PAMPs) in the fungal cell wall – specifically recognition of the β 1,3-glucan and the chitin moieties found in the inner cell wall (*29-32*). However, recently it has been shown that *C. albicans* have adapted to physically mask PAMPs such as chitin from the receptors (*33*). As chitin is demonstrated to be essential to hyphal growth and sustained virulence, development of methods to study the molecular integration and distribution of this polymer in the cell wall is essential.

In this work, a strategy to embed bio-orthogonal probes into chitin is developed. This is substantiated by the use of various appropriate knockouts strains, microscopy, mass spectrometry and flow cytometry analysis. Importantly, the budding to hyphae switch that enables *C. albicans* pathogenesis is not disturbed by the probes. This method will be a powerful tool for the community to study the mechanism of fundamental chitin synthases/degraders across the animal kingdom, identify new antifungal regimes, and determine new methods to modulate the human immune system.

### Modified GlcNAc fragments are accepted into the chitin biosynthesis pathway via Hxk1

To label the chitin backbone, modified GlcNAc fragments were obtained. A library of GlcNAc derivatives were prepared including: (**1**) Azide, (**2**) Alkyne, (**3**) Tetrazine, (**4**) NBD, (**5**) and peracetylated azide (Fig. S2). These handles were all installed at the 2-N position of GlcNAc (NAG). The first three include standard click modalities, with (**1**-**3**) being demonstrated as handles capable of being incorporated into the PG layer of the bacterial cell wall when positioned on the molecule *N*-acetylmuramic acid (MurNAc) via the use of copper-catalyzed azide-alkyne cycloaddition (CuAAc) (*17, 34*). Whereas probes **1** and **2** have also been used successfully in *S. cerevisiae* by Bertozzi and co-workers (*18*). **3** includes a tetrazine modality – the tetrazine–*trans*-cyclooctene ligation (Tz-TCO) is a rapid, non-cytotoxic bio-orthogonal reaction to cells that has been frequently used in visualization of a variety of biological processes (*35*). Recently, we have shown that this minimalist tetrazine probe, H-Tz, is capable of incorporation into bacterial PG (*34*) and hypothesized that a similar labeling strategy could be applied to filamentous yeast to rapidly label the chitin layer of the fungal cell wall. The NBD probe, **4**, initially developed by Vocadlo to label GlcNAc modified proteins (*36*), enables direct fluorescent modification of the polymer negating the need of a “click” reaction. Finally, peracetylated probe, **5**, is commercially available and was used here to further determine the mechanism of uptake of the GlcNAc probes and their incorporation into chitin.

It is hypothesized that if exogenous GlcNAc is incorporated in the cell wall, the kinase Hxk1 from *C. albicans* is involved. Initially characterized in 2001, Hxk1 has been shown by thin layer chromatography (TLC) to phosphorylate sugars such as GlcNAc, glucose, and mannose (*8*). This substrate promiscuity differs from the enzyme Hxk2, which shows restricted substrate usage, capable of only phosphorylating hexose; this enzyme found in both *C. albicans* and *S. cerevisiae* is responsible for phosphorylating glucose which can be further incorporated into chitin (Fig. 1) (*6, 37, 38*). In order to assess Hxk1 substrate specificity, the enzyme was expressed and purified (Fig. S1) and assessed using an *in vitro* biochemical mass spectrometry-chemoenzymatic assay. In agreement with previous reports (*8*), the data show that purified Hxk1 is capable of accepting GlcNAc and mannose, producing phosphorylated products (Fig 2A) (Table S1). In addition, the data show that the GlcNAc based probes, (**1-4**), are accepted and phosphorylated by Hxk1. (Fig. 2A) (Table S1). However, the enzyme does not have activity with MurNAc, the other carbohydrate component of bacterial peptidoglycan (*39*) (Fig. 2A) (Table S1) and probe (**5**), per acetylated GlcNAc, was not assessed, as it does not have a free hydroxy to phosphorylate.

**Fig. 2.**
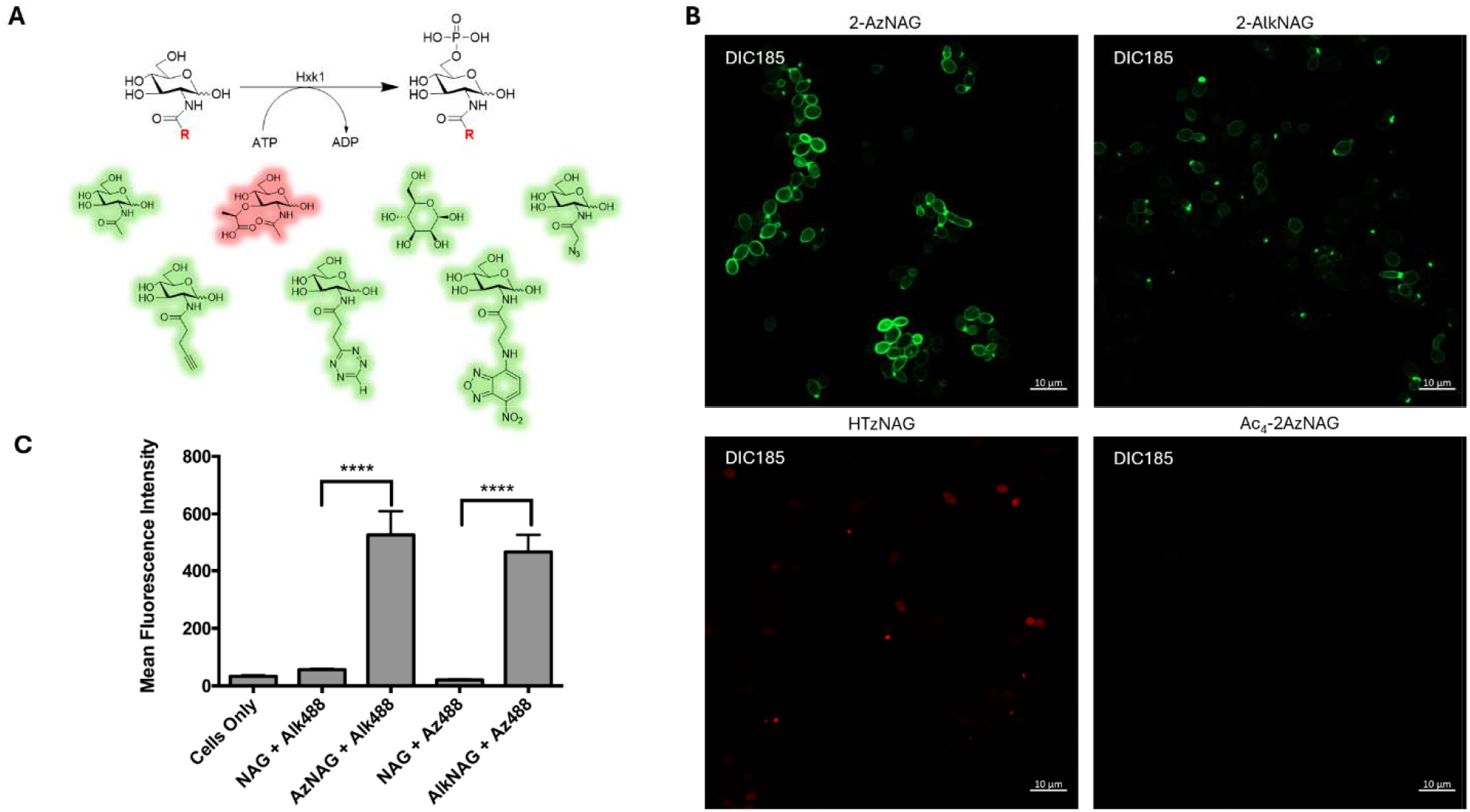
NAG probe incorporation into *C. albicans* WT. (**A**) Chemoenzymatic screen of Hxk1 activity on various sugars. Samples were subjected to high-resolution LCMS, ESI negative mode. (**B**) Confocal fluorescent images of CuAAC labeled *C. albicans* WT with NAG based probes in fixed conditions. Scale bars, 10 μm (**C**) MFI values of *C. albicans* WT cells when labeled in fixed conditions with either AzNAG or AlkNAG. Statistics were analyzed using GraphPad Prism 7 via a 2 way ANOVA test.

Konopka and co-worker have previously reported that *C. albicans* can grow on GlcNAc as a carbon source when restricted (i.e. the only carbon source) (*6, 40, 41*). Here, growth conditions in which *C. albicans* would not need to use GlcNAc as a carbon source were chosen, potentially permitting the cell to selectively shuttle the NAG probe to the chitin polymer; thereby restricting the distribution of the probe. Following a previously published protocol (*42*), when probes **1** and **2** were incubated in mid log phase for 2 hours, the cell wall was robustly labeled (Fig. 2B, Fig. S3-4). Qualitatively, more robust labeling was observed with the 2-AzNAG probe than the alkyne; however, this observation could be due to the alkyne handle being reduced in the acidic growth media. To assess labeling efficiency, flow cytometry was used to show that both the AzNAG and the AlkNAG probes are incorporated into the cell compared to the NAG and cells only control. However, the data show that the AzNAG probe labeled a higher number of cells compared to that of the AlkNAG probe (Fig. 2C, Fig. S10-11). Alternatively, incubation with the tetrazine probe (**3**) did not show clear labeling of the cells, despite the probe being phosphorylated by Hxk1 (Fig. 2B), this could be a result of the fact that tetrazines are extremely acid sensitive. Incubation with the NBD probe (**4**) also did not show labeling of cells (Data not shown) which could be due to size or chemical properties leading to the probe not being accepted by the cell. Importantly the acetylated probe (**5**) did not label the cell wall (Fig. 2B, Fig. S5), implying that *C. albicans* lacks cellular deacetylases that can process and unmask this probe or that the specific GlcNAc transporter does not accept this probe.

### Chitinase digestion indicates probe incorporation into mature chitin

Incorporation of the probes into the chitin moiety of the cell wall of *C. albicans* was determined using high-resolution mass spectroscopy (HRMS). Cells treated with either NAG or AzNAG were lysed, and sugars including chitin were isolated. Chitinase cocktails were utilized to digest chitin in isolated samples. Disaccharide and tetra-saccharide subunits of chitin were identified in the AzNAG treated samples (Fig. 3) and in the digested chitin control sample (Fig. S12-13), indicating that AzNAG is incorporated into the chitin polymer.

**Fig. 3.**
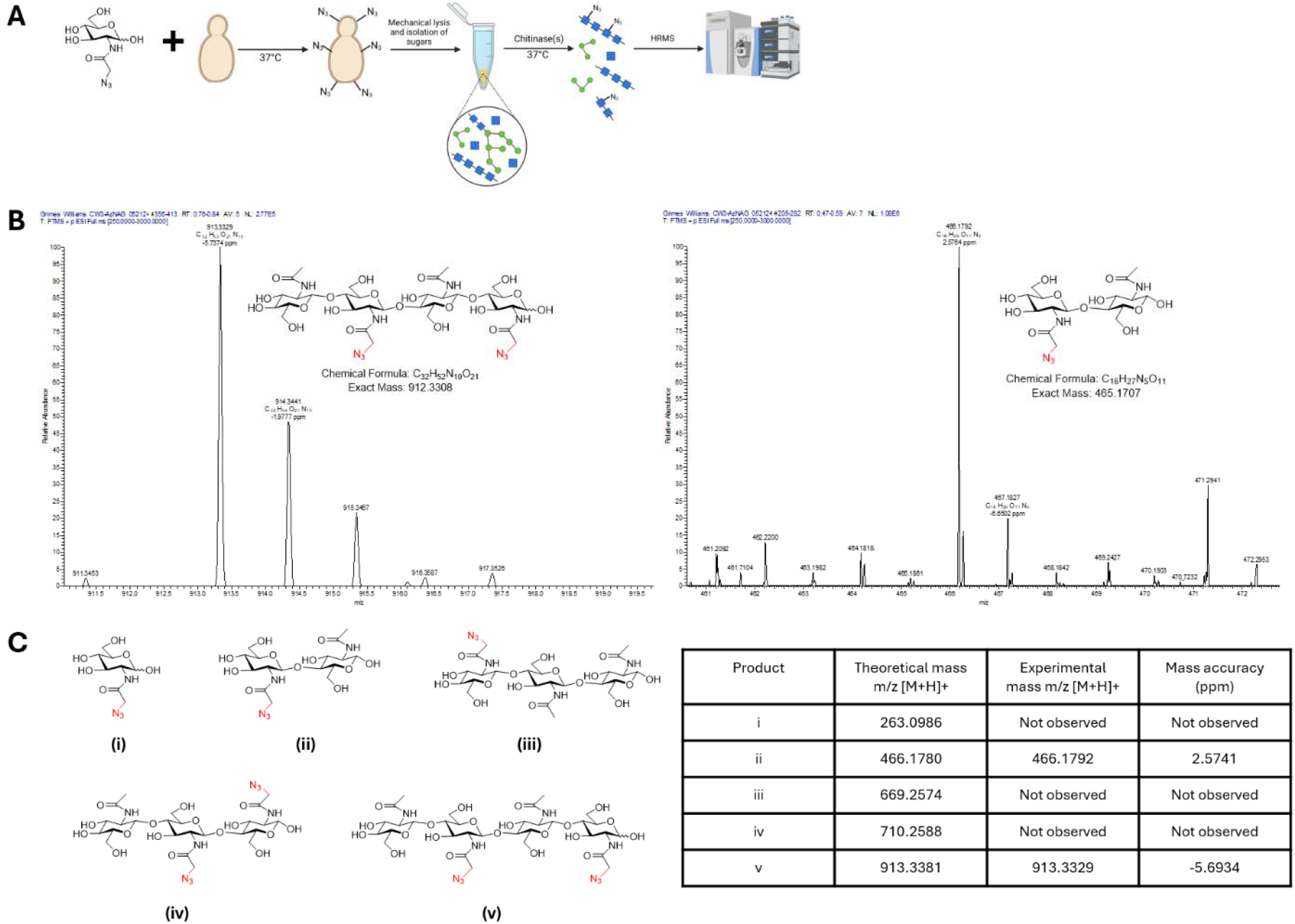
Mass spectrometry-based analysis of NAG probe incorporation into cells. (**A**) Workflow for determining AzNAG incorporation into the chitin polymer in *C. albicans* cells. (**B** Chitinase digestion of AzNAG labeled chitin; analyzed via HRMS for disaccharide and tetra-saccharide fragments in ESI positive mode. (**C**) Chemical structures of fragments analyzed in HRMS studies; table of expected mass, observed mass, and mass accuracy (ppm) of chitin fragments.

To confirm that the probes are going through the proposed chitin biosynthesis pathway (Fig. 1), yeast cell wall labeling was attempted in cells in which the transporter NGT1, a specific GlcNAc transporter (*40*), is knocked out (strain YJA3). If the NAG probe is recognized and transported by the dedicated GlcNAc transporter, Ngt1 this would suggest that the carbohydrate is transported across the membrane and does not passively diffuse. Attempts to label YJA3 with AzNAG were successful (Fig. 4A, Fig. S6), indicating that there could be an alternative transporter besides NGT1 that allows for GlcNAc transport into the cell when NGT1 is no longer present. Alternatively, when the genes of the GlcNAc catabolism enzymes Hxk1, Nag1, and Dac1 (*6*) are knocked out (strain AG738), incubation with the AzNAG probe did not show incorporation into the fungal cell wall (Fig. 4D, Fig. S7). This indicates that Hxk1 is essential and likely required for GlcNAc incorporation into the fungal cell wall.

**Fig. 4.**
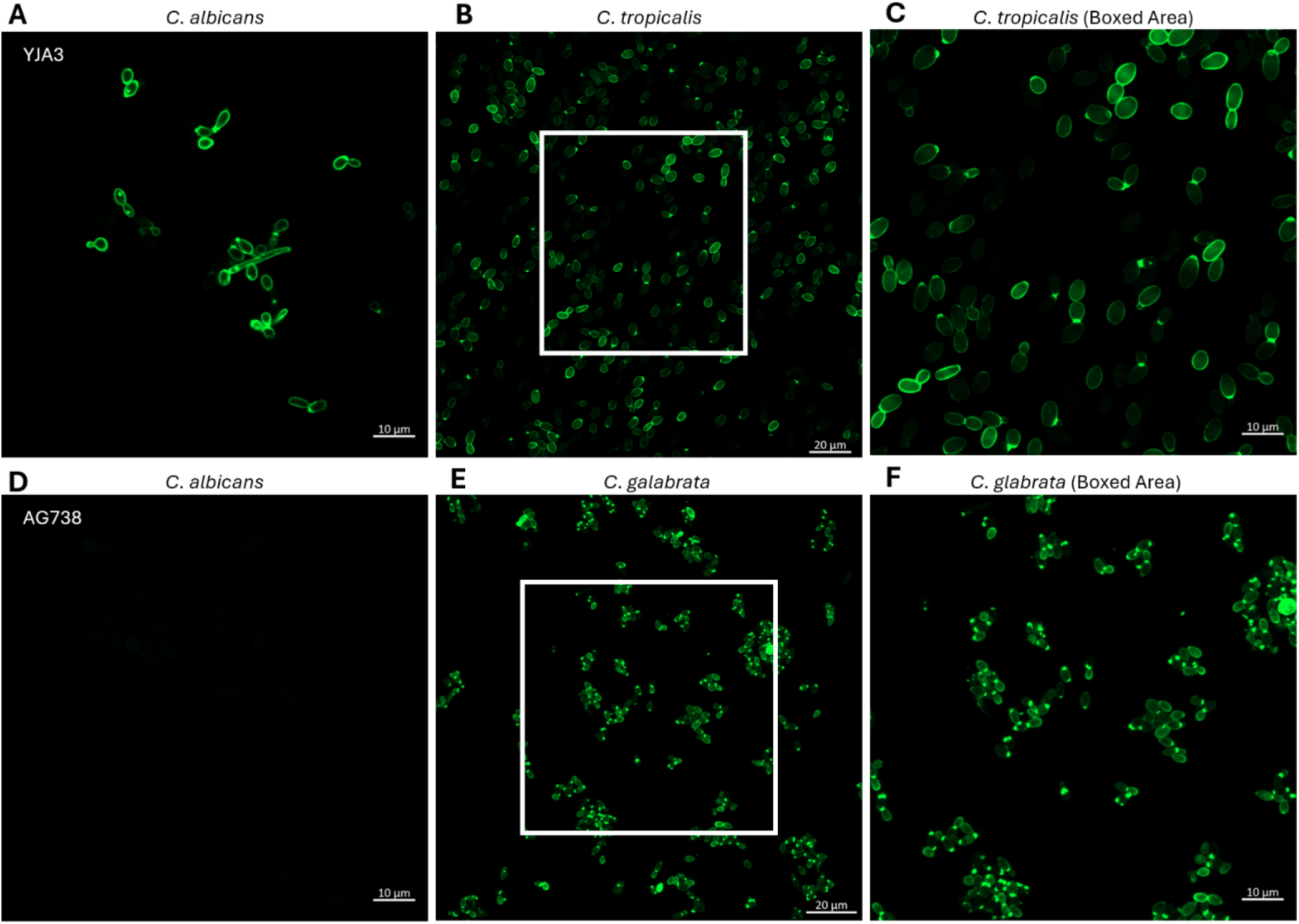
Labeling of alternative Candida strains with 2-AzNAG probe. (**A**) C. albicans knockout strain YJA3, where the GlcNAc specific transporter NGT1 is knocked out. (**B**) *C. tropicalis* WT. (**C**) *C. tropicalis* zoomed in using a 63x lens. (**D**) C. albicans knockout strain AG738, where three GlcNAc catabolism genes are knocked out. (**E**) *C. glabrata* WT. (**F**) *C. glabrata* zoomed in using a 63x lens.

### NAG probes are incorporated into chitin in alternative pathogenic Candida species

Although *C. albicans* is the most common fungal pathogen to cause candidiasis in humans, four other Candida species are also recognized as significant contributors to the development of candidiasis: *Candida glabrata* (*C. glabrata*), *Candida tropicalis* (*C. tropicalis*), *Candida parapsilosis* (*C. parapsilosis*), and *Candida krusei* (*C. krusei*) (*43, 44*). A search for homologs of Hxk1 from *C. albicans* in these other Candida species indicated that a variety of filamentous Candida strains do have a homolog of this enzyme. We tested *C. tropicalis* (67.6% sequence homology) ability to label with the AzNAG probe. *C. tropicalis* incorporates this probe into its fungal cell wall (Fig. 4B and 4C, Fig. S8), indicating that the use of these novel probes is not isolated to *C. albicans* but likely all species that contain a homolog of Hxk1 that can phosphorylate GlcNAc and incorporate downstream products into its fungal cell wall. To support this hypothesis, *C. glabrata* was shown to also robustly label with the AzNAG probe (Fig. 4E and 4F, Fig. S9) despite only having 23.8% sequence homology.

### Filamentous cells sense and incorporate NAG probes into the hyphal cell wall

The probe was assessed for its ability to integrate into the cell wall when polymorphic fungi, such as *C. albicans*, are making the transition into their filamentous hyphal form (Fig. 1). Upon incubation with the AzNAG or the AlkNAG probe and a known hyphal inducer, robust hyphal labeling was observed, indicating that the probes do not interfere with hyphal growth (Fig. 5A).

**Fig. 5.**
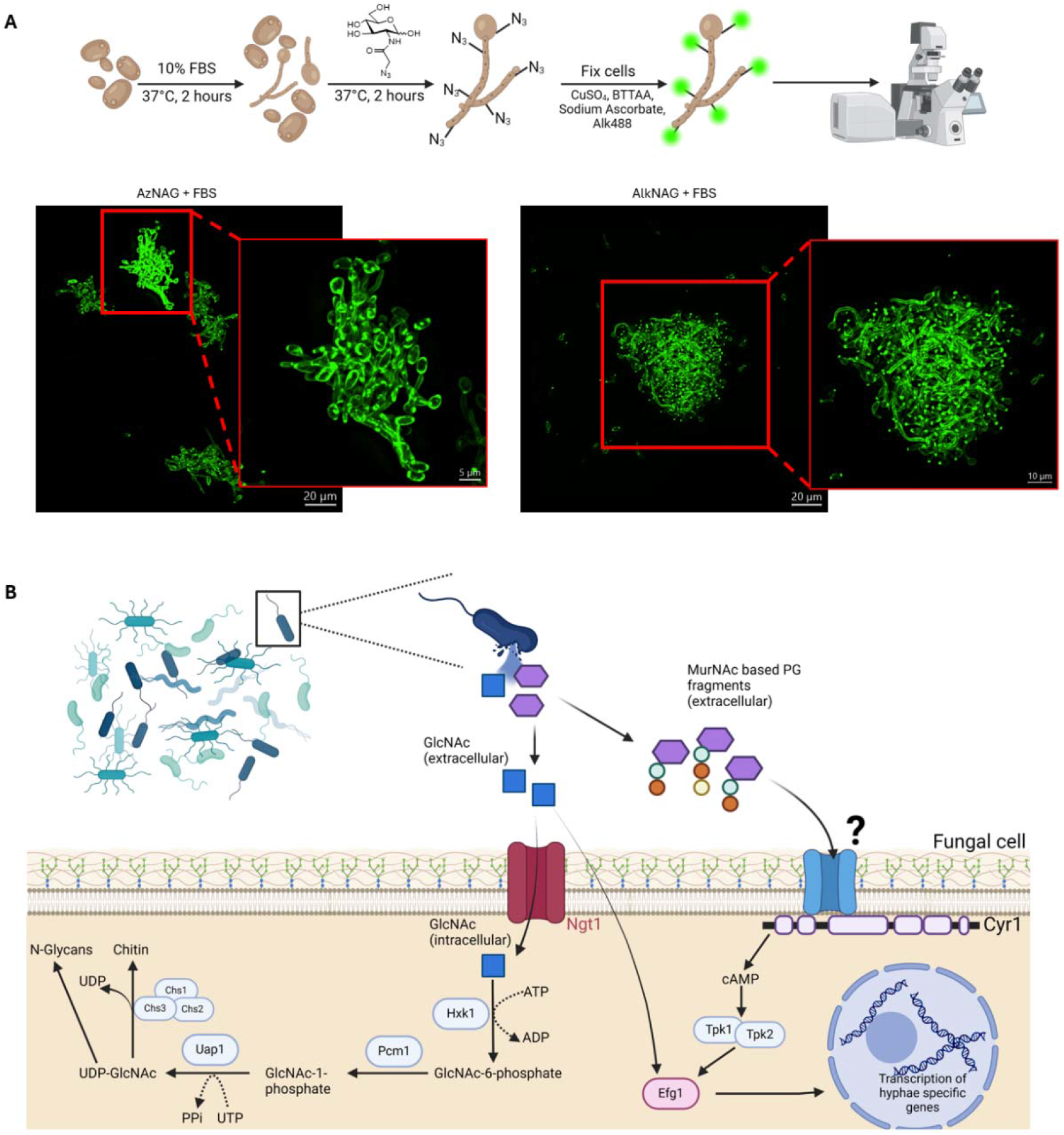
Incorporation of NAG probes in hyphae morphology of *C. albicans* WT cells. (**A**) Workflow for labeling of *C. albicans* WT hyphal cells with NAG probes. *C. albicans* WT hyphal cells visualization with either AzNAG or AlkNAG under fixed conditions. Deconvolution achieved through Imaris software. (**B**) *C. albicans* multifaceted use of PG fragments shed from bacteria.

Here the hyphal conditions were triggered with the known hyphal inducer fetal bovine serum (FBS). FBS contains numerous proteins and nutrients that contribute to the induction of this morphogenic transition. Interestingly, the mixture also contains bacterial cell PG fragments (*45*). Bacterial PG is a polymer consisting of alternating residues of β-1,4 linked GlcNAc and MurNAc (*39*). In 2008, Wang and co-workers showed that discrete peptidoglycan fragments that contain MurNAc, such as muramyl dipeptide (MDP), was the prominent PG fragment in serum that induced hyphal growth (*45*). It is hypothesized that PG and other bacterial derived molecules in serum induce hyphal formation by binding directly to Cyr1p in *C. albicans* (Fig. 5B). Binding of fragments to Cyr1p will increase intracellular cAMP levels, subsequently activating the PKA pathway, and triggering hyphal growth (*46-48*).

Here the data show that when cells are co-treated with the AzideNAG probe and PG fragments of FBS, robust hyphae labeling is observed (Fig. 5A). These data suggest that the bacteria-fungus relationship is complex and that the *C. albicans* are able to use peptidoglycan in a multifaceted manner: 1. to sense the presence of bacteria and then 2. To take advantage of the released GlcNAc to grow the chitin cell wall. It is known that when *C. albicans* transforms from budding to hyphae, the amount of chitin needed is approximately 3x higher than in budding cells (*26, 49*). Thus, the yeast are opportunistic, taking advantage of the GlcNAc from the PG to use as a new cell wall building block. The other PG building block, MurNAc, serves a sensor function to permit this transition to the organism’s pathogenic form as certain MurNAc containing PG fragments induce hyphae growth in different quantities. (Fig. 5B) (Fig. S14-15). To assess this, it is shown that when cells are co-treated with NAG and commercially available MDP, robust hyphal growth is observed at earlier time points than when cells are treated with MDP alone (Fig. S16), suggesting a dual use of the bacterial peptidoglycan by the pathogenic yeast.

## Discussion

A robust method to label the chitin of wild-type, pathogenic *C. albicans* has been developed. This method allows for continuous growth of chitin in the inner layer of the fungal cell wall, contrary to current labeling methods that utilize Calcofluor white (*13*). Robust labeling was observed with either an azide or an alkyne handle and these handles do not affect the cell’s ability to transform into its filamentous, pathogenic state; however protected probes do not have the ability to be incorporated into the cell, as the cell likely lacks the machinery to process these carbohydrate moieties. Unlike the model organism *S. cerevisiae*, which lacks the specific kinase Hxk1, filamentous fungi that contain a homolog of the enzyme Hxk1 can also be labeled with this method as observed with *C. tropicalis* and *C. glabrata*.

The method has permitted the identification of a two-pronged strategy that the pathogen uses to sense the presence of bacteria and transform from a non-pathogenic state into the pathogenic hyphae form. This work suggests that inhibiting GlcNAc uptake and incorporation in chitin biosynthesis will serve as excellent targets for anti-fungal strategies. This method will enable the study of the fundamental chitin synthases/degraders by providing new tools and labeled substrates. Future work will determine if these GlcNAc probes can be used to label chitin across the animal kingdom.

## Acknowledgments

Research reported in this publication was supported via instrument and centers (NMR, Microscopy) by National Institute of General Medical Sciences or the National Institutes of Health under Award Numbers P20GM104316, P20GM103446, S10OD030321, and P20GM139760. We would like to thank Jeff Caplan and the Delaware Biotechnology Institute Bioimaging Center, William Trout; Shi Bai and the UD NMR facility; and PapaNii Asare-Okai, Katherine Martin, and Elijah Hudson from the UD Mass Spectrometry Core Facility. We would also like to thank James Konopka (Stony Brook University) for supplying us with various strains of *C. albicans* and the Blenner Lab at the University of Delaware for providing us with *C. tropicalis* and *C. glabrata* strains in our studies. Figures were created using BioRender.

## Funding

National Institutes of Health grant T32GM133395 (CW, SNH)

Camille and Henry Dreyfus Foundation (CW)

P01AI172525 (BRC)

## Author contributions

Conceptualization: CW, BRC, and CLG

Formal Analysis: CW and SNH

Methodology: CW

Investigation: CW and BRC

Visualization: CW

Writing – original draft: CW and CLG

Writing – review & editing: CW, BRC, SNH, and CLG

## Data and materials availability

A full description of the methods can be found in the supplementary materials. This includes labeling protocols, cell-based imaging protocols, and synthesis of all compounds. We can provide small amounts of NAG samples if available; however, the rigorous protocols provided in SI will allow for the production in any lab; we are open to helping labs troubleshoot the synthesis if necessary by contacting.

## Supplementary Materials

Materials and Methods

Figs. S1 to S24

Tables S1

References (*8, 34, 42, 50–52*)

MDAR Reproducibility Checklist

