## Supporting Information for "Bioorthogonal labeling of chitin in pathogenic Candida species reveals biochemical mechanisms of hyphal growth and homeostasis"

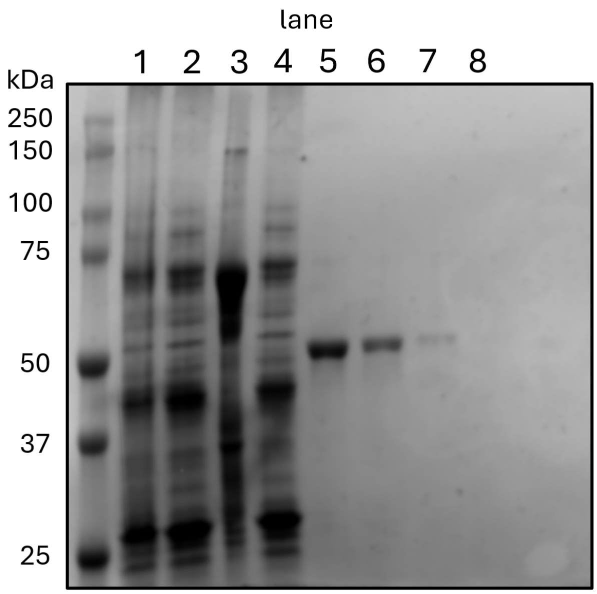


**Fig. S1. Purification of Hxk1.** Coomassie stained gel of purified Hxk1 run on a 10% TGX polyacrylamide gel. Lane samples – 1: lysate, 2: supernatant, 3: pellet, 4: flowthrough, 5: elution 1, 6: elution 2, 7: elution 3, 8: elution 4.

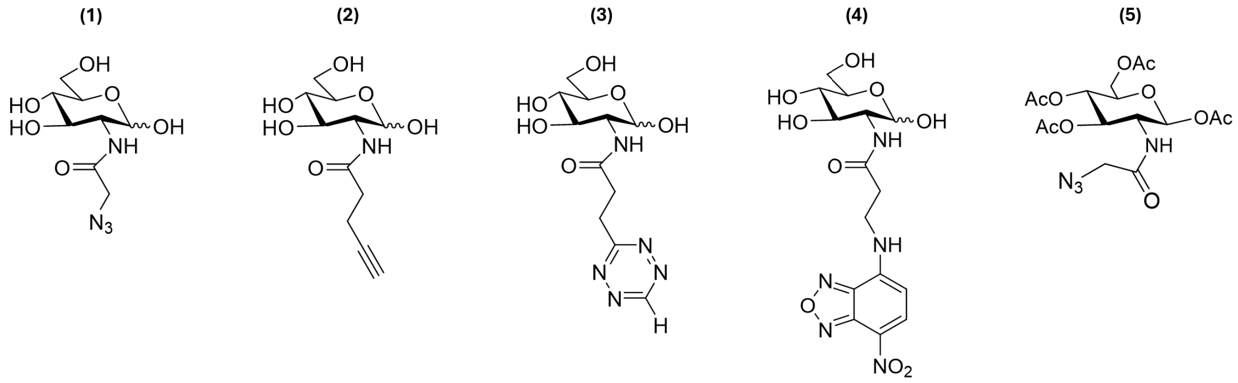


**Fig. S2. Chemical structures of NAG probes used to bio-orthogonally label *C. albicans*.** (1) AzNAG (2) AlkNAG (3) HTzNAG (4) NBD NAG (5) $Ac_4$AzNAG. $Ac_4$AzNAG was purchased from Click Chemistry Tools (acquired by Vector Laboratories). AzNAG, AlkNAG, HTzNAG, and NBD NAG were synthesized in our lab as described above (reference section above about synthesis).

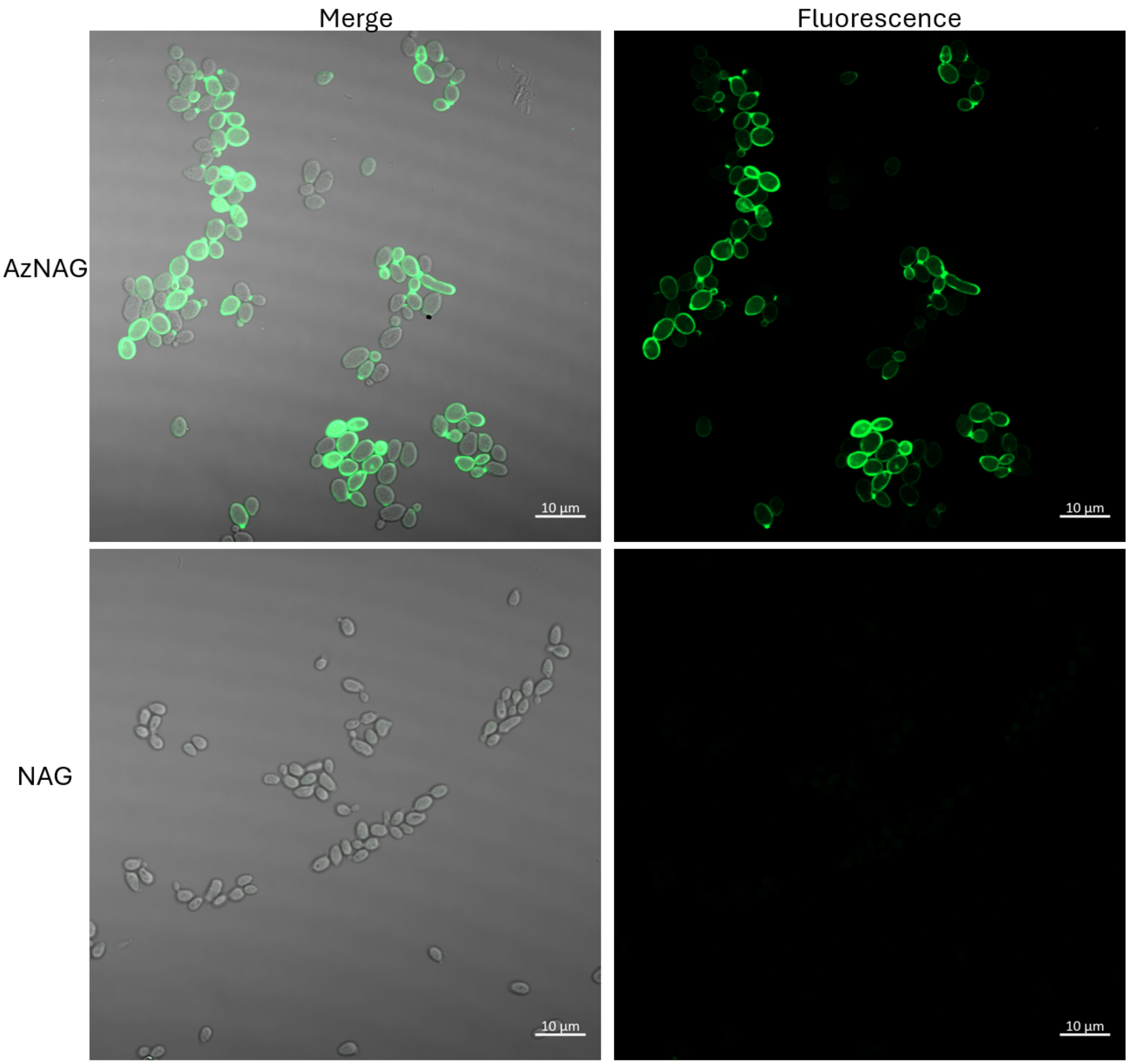


**Fig. S3. CUAAC labeling of AzNAG in *C. albicans*.** Merged (left) Brightfield and Fluorescent (right) images of DIC185 cells treated with 3mM AzNAG or NAG and clicked with 20 µM Alk488 (green) (scale bars, 10 µm). Images are representative of a minimum of three fields viewed and the experiment was conducted in at least three biological replicates.

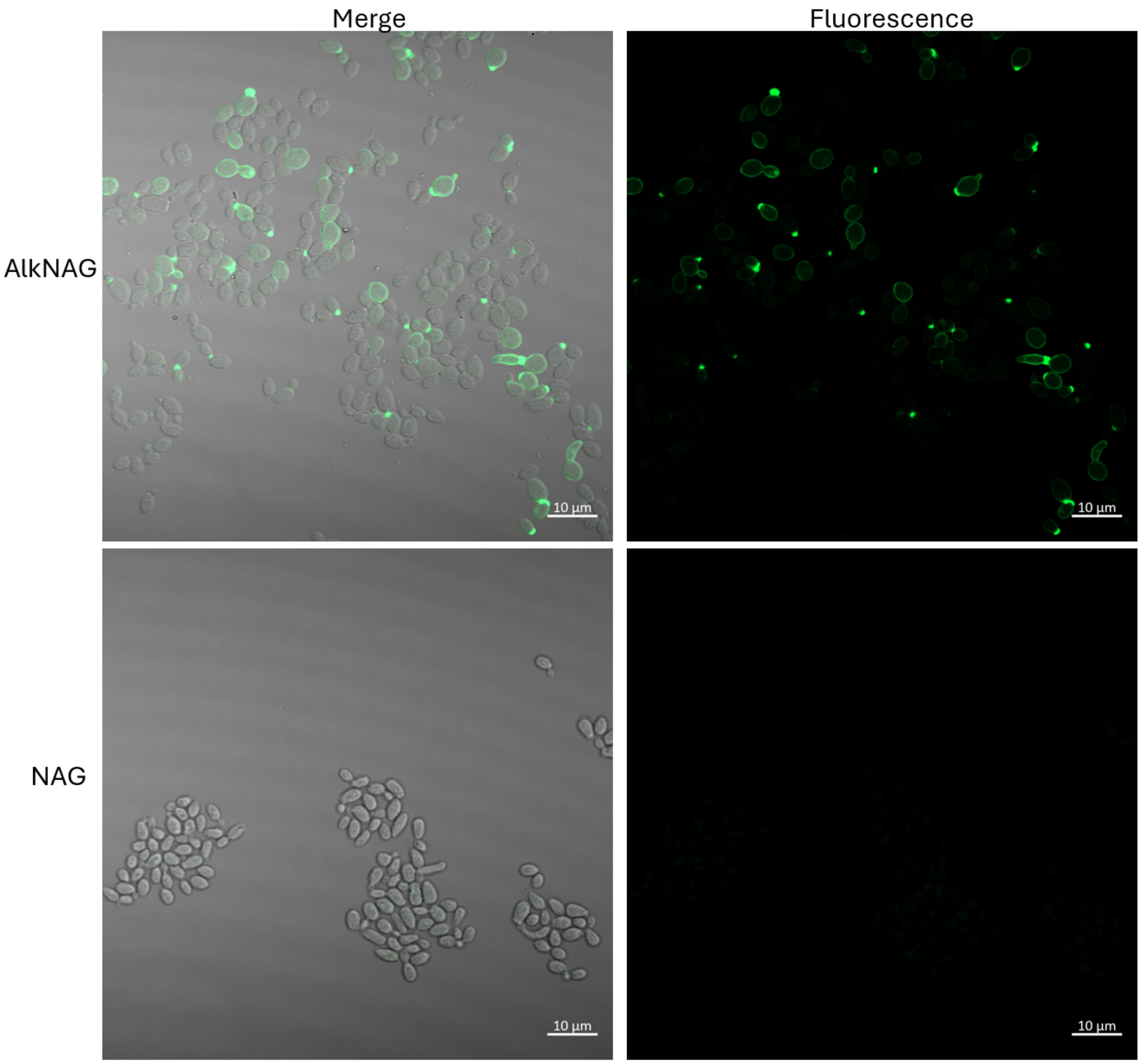


**Fig. S4. CUAAC labeling of AlkNAG in *C. albicans*.** Merged (left) Brightfield and Fluorescent (right) images of DIC185 cells treated with 3mM AlkNAG or NAG and clicked with 20 µM Az488 (green) (scale bars, 10 µm). Images are representative of a minimum of three fields viewed and the experiment was conducted in three biological replicates.

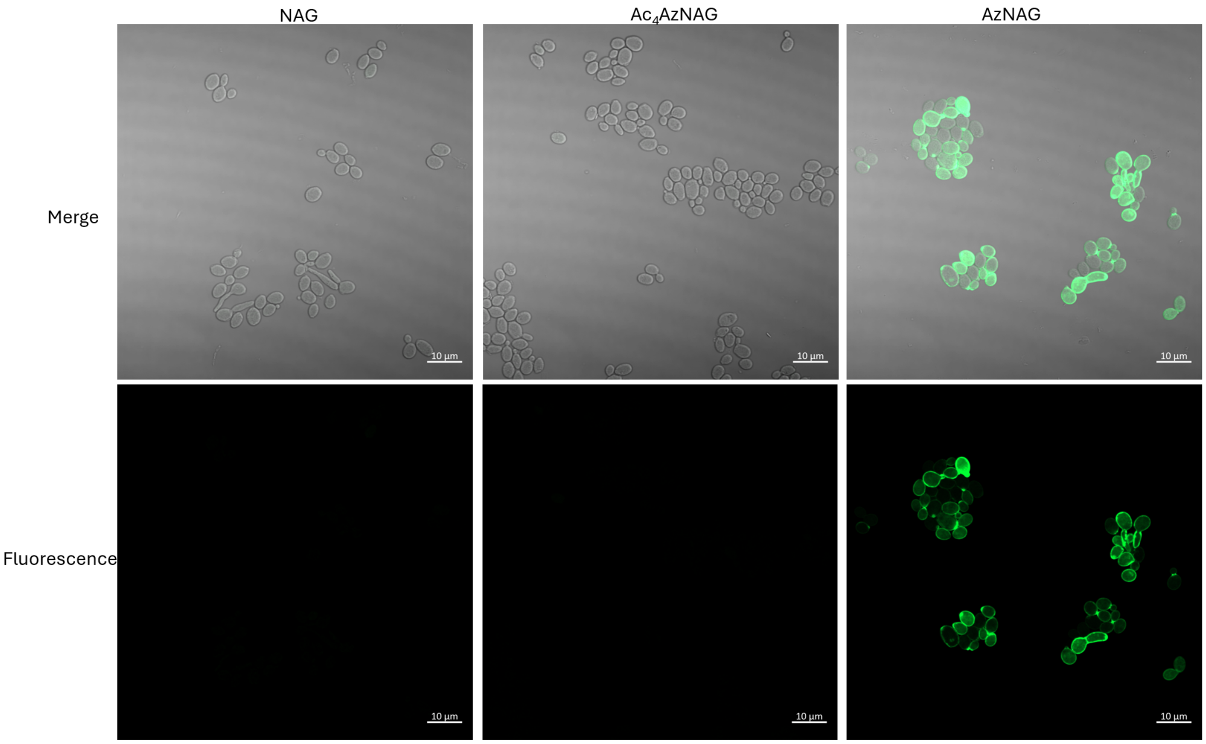


**Fig. S5. CUAAC labeling of $Ac_4AzNAG$ in *C. albicans*.** Merged (top) Brightfield and Fluorescent (bottom) images of DIC185 cells treated with 3mM NAG, $Ac_4AzNAG$, or NAG and clicked with 20 µM Alk488 (green) (scale bars, 10 µm). Images are representative of a minimum of three fields viewed and the experiment was conducted in three biological replicates.

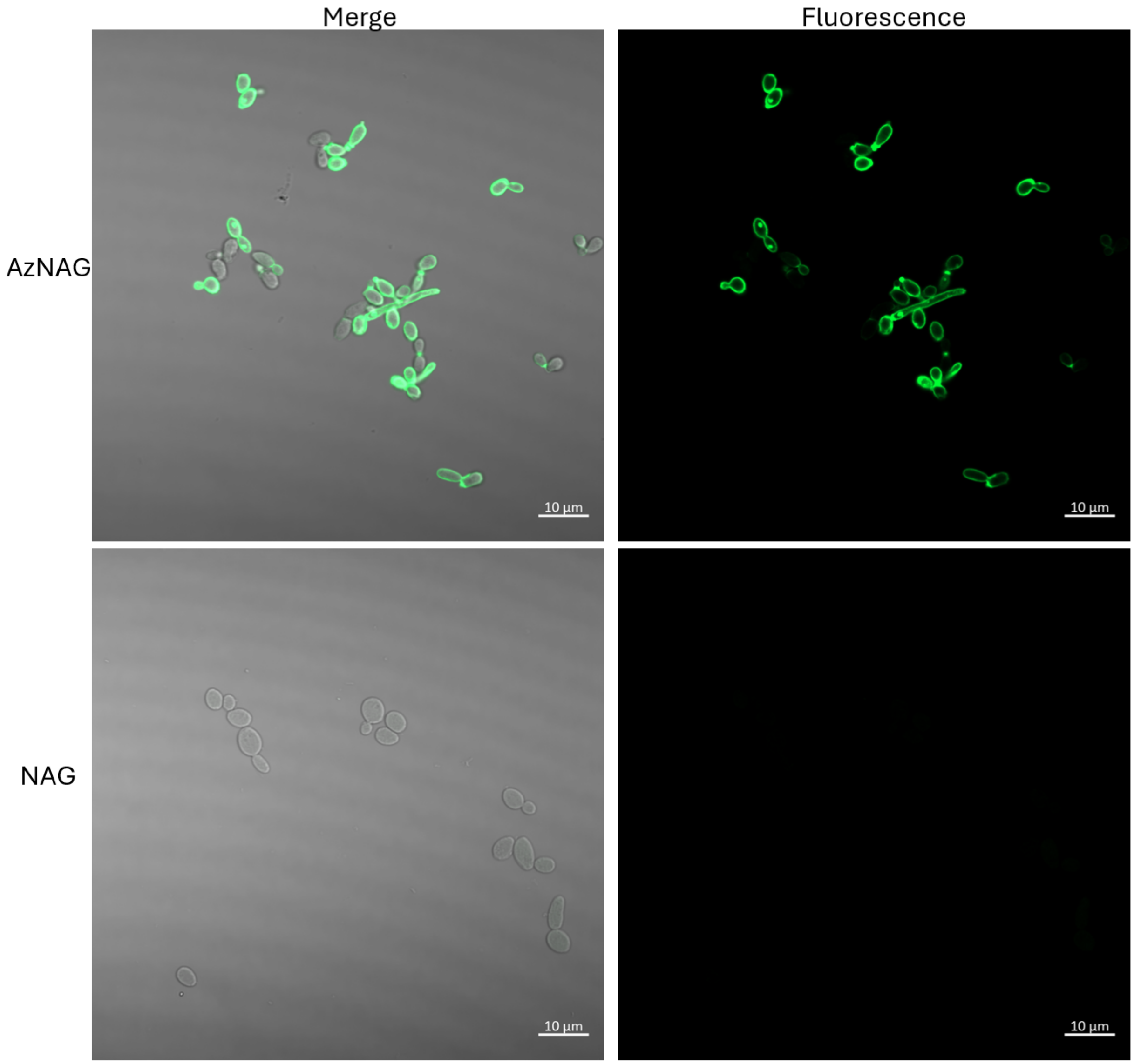


**Fig. S6. CUAAC labeling of AzNAG in *C. albicans* strain YJA3.** Merged (left) Brightfield and Fluorescent (right) images of YJA3 cells treated with 3mM AzNAG or NAG and clicked with 20 µM Alk488 (green) (scale bars, 10 µm). Images are representative of a minimum of three fields viewed and the experiment was conducted in three biological replicates.

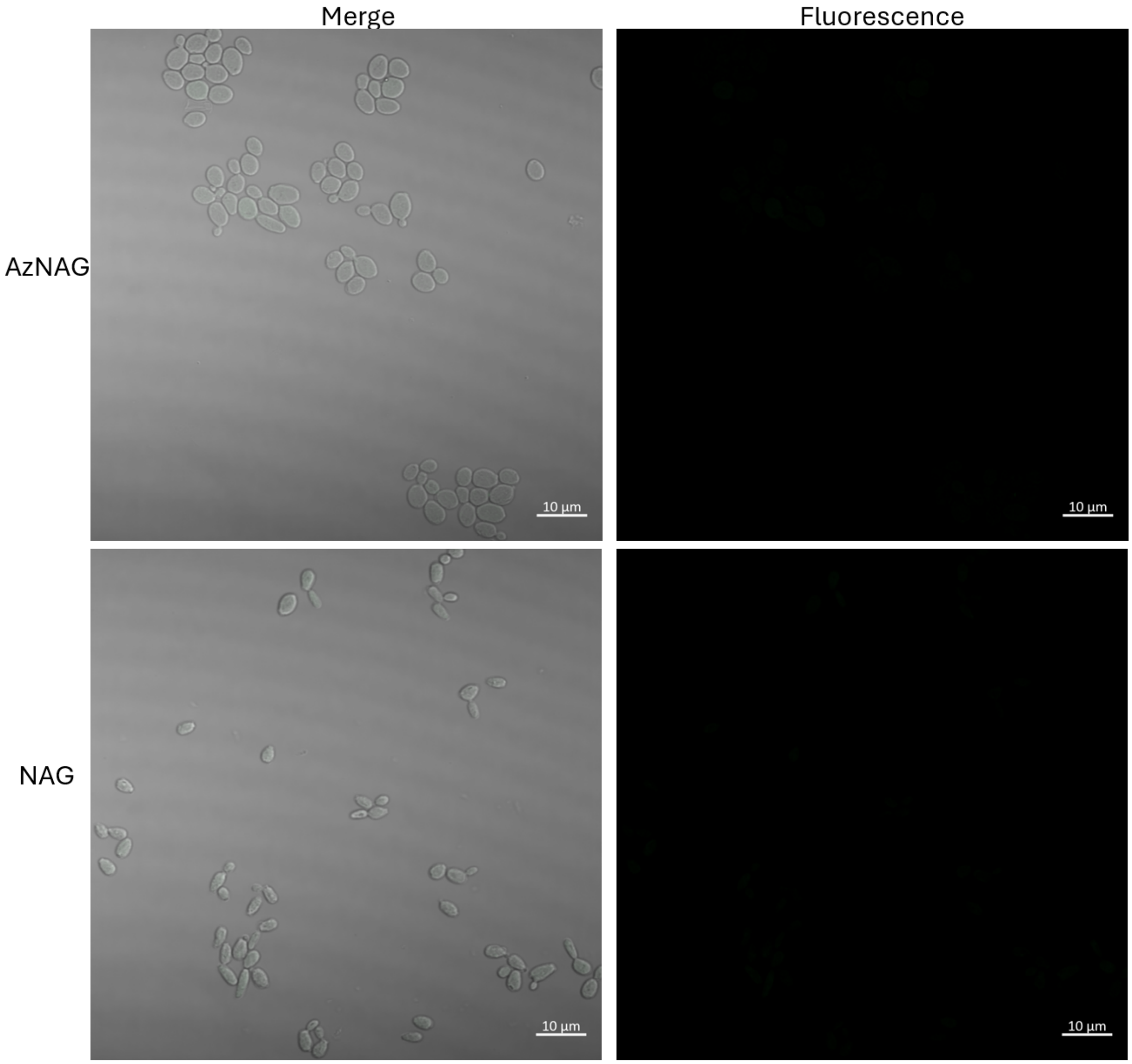


**Fig. S7. CUAAC labeling of AzNAG in *C. albicans* strain AG738.** Merged (left) Brightfield and Fluorescent (right) images of AG738 cells treated with 3mM AzNAG or NAG and clicked with 20 µM Alk488 (green) (scale bars, 10 µm). Images are representative of a minimum of three fields viewed and the experiment was conducted in three biological replicates.

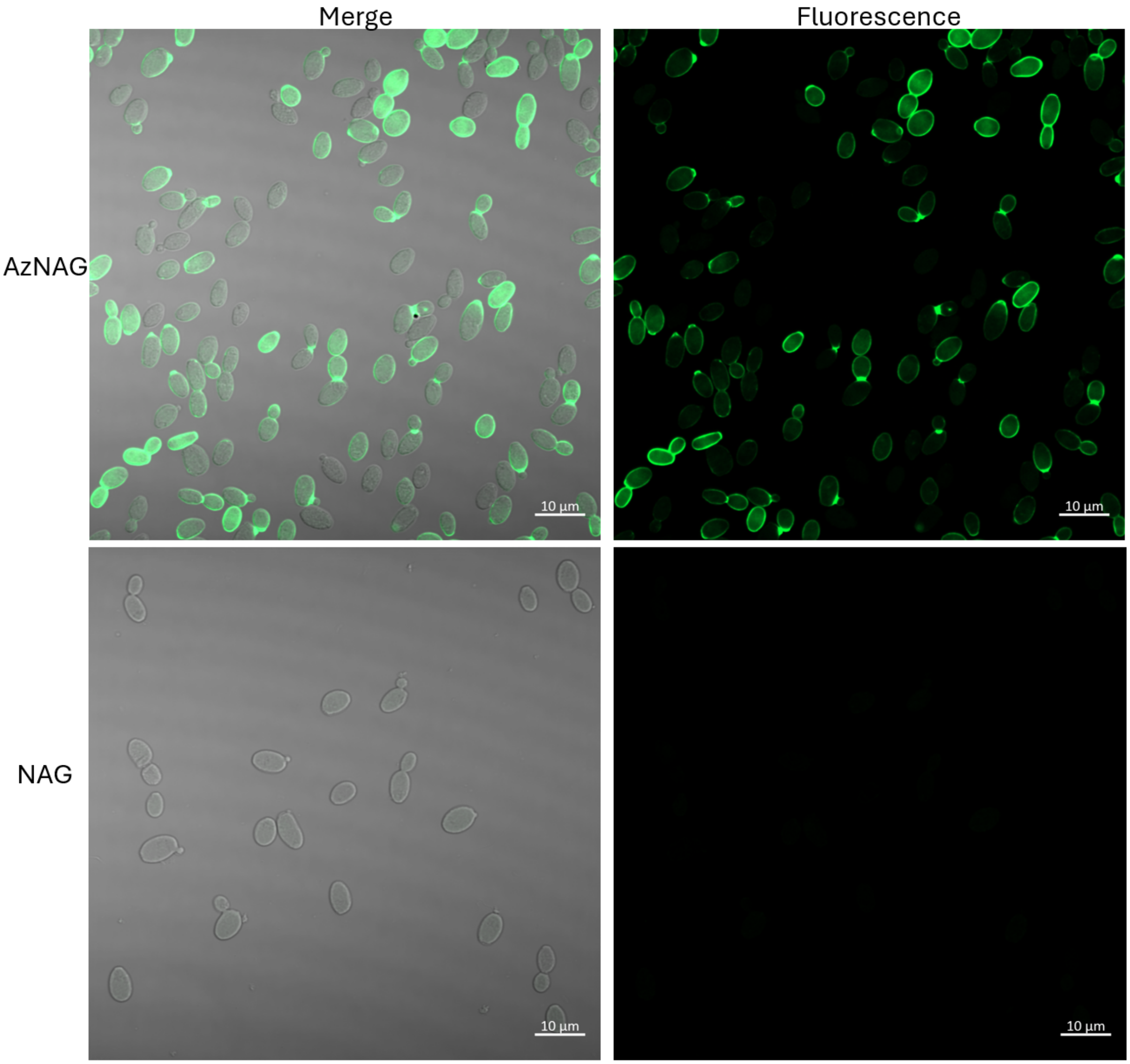


**Fig. S8. CUAAC labeling of AzNAG in *C. tropicalis*.** Merged (left) Brightfield and Fluorescent (right) images of *C. tropicalis* cells treated with 3mM AzNAG or NAG and clicked with 20 µM Alk488 (green) (scale bars, 10 µm). Images are representative of a minimum of three fields viewed and the experiment was conducted in three biological replicates.

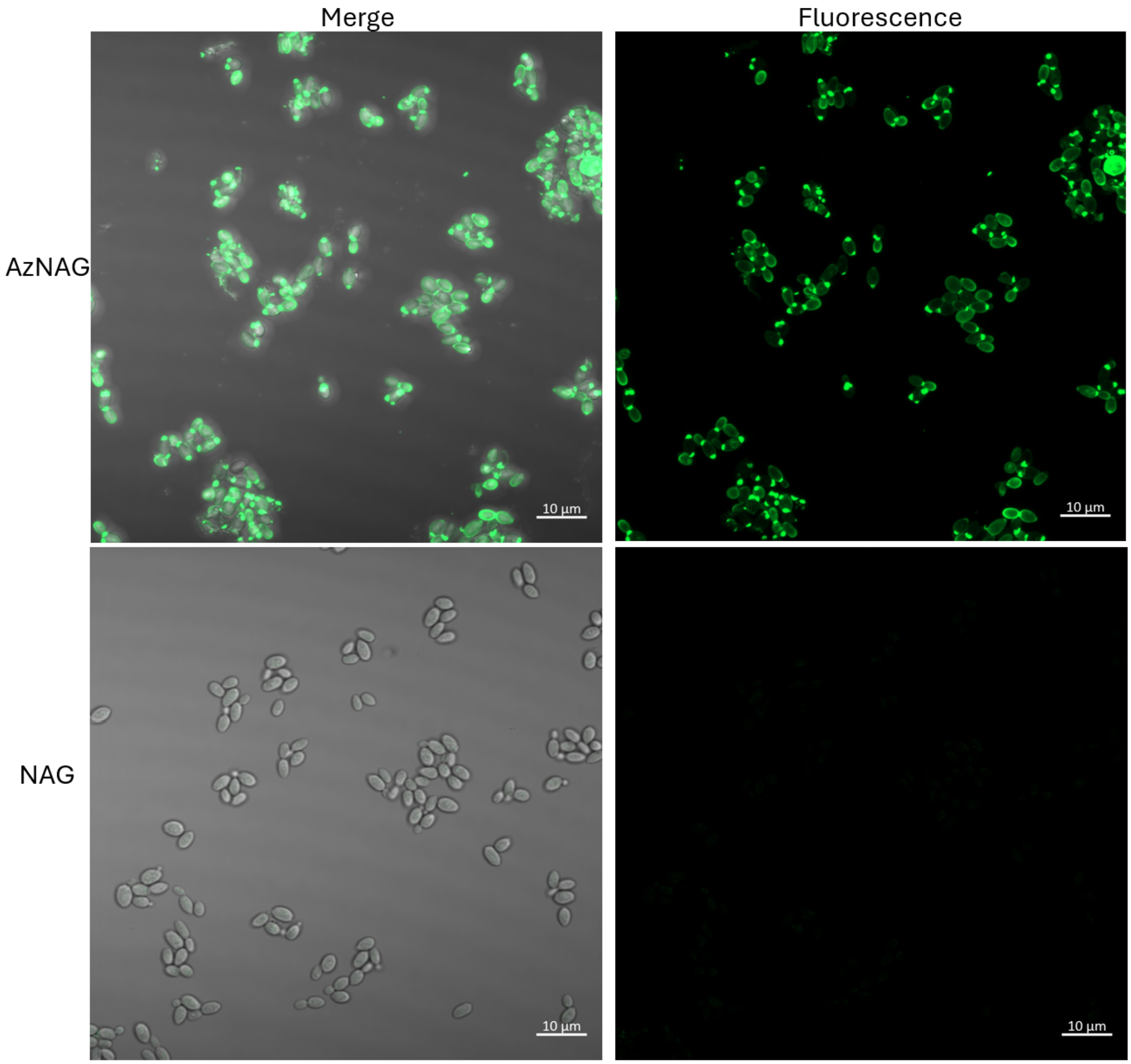


**Fig. S9. CUAAC labeling of AzNAG in *C. glabrata*.** Merged (left) Brightfield and Fluorescent (right) images of *C. glabrata* cells treated with 3mM AzNAG or NAG and clicked with 20 µM Alk488 (green) (scale bars, 10 µm). Images are representative of a minimum of three fields viewed and the experiment was conducted in three biological replicates.

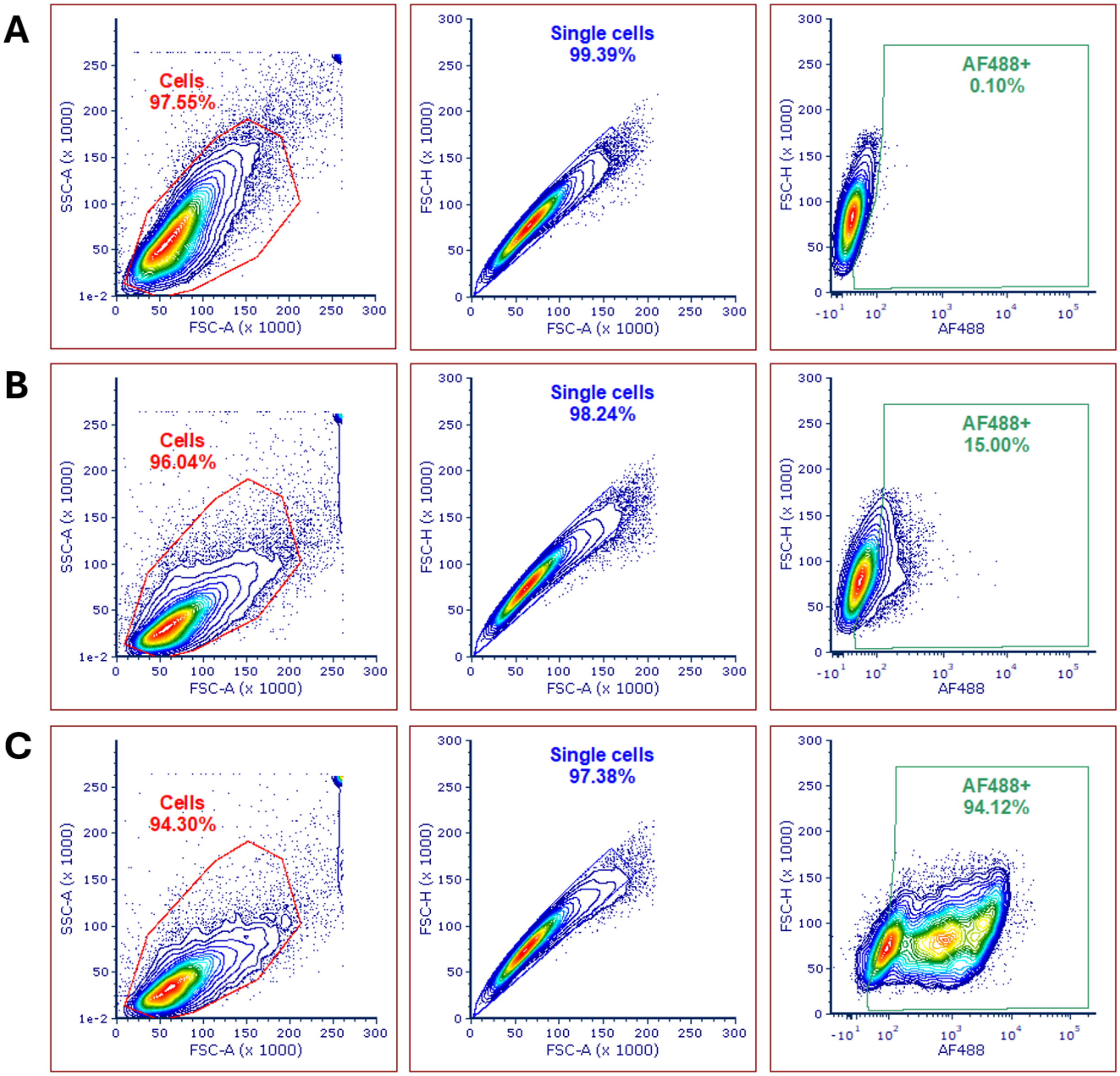


**Fig. S10. Flow cytometry population comparison of AzNAG labeling in *C. albicans* WT.** DIC185 cells only (A) or DIC185 cells remodeled with 3mM NAG (B) or AzNAG (C) and then clicked with 20 µM Alk488. Flow samples were processed using a BD FACS Aria Fusion Cell Sorter using 50,000 cell counts. Data was processed in FCSExpress 7. The gating method was created using 0.1% TRITC channel signal in the cells only sample and then applied to the experimental samples. Data shown is representative of three biological replicates from at least two technical replicates.

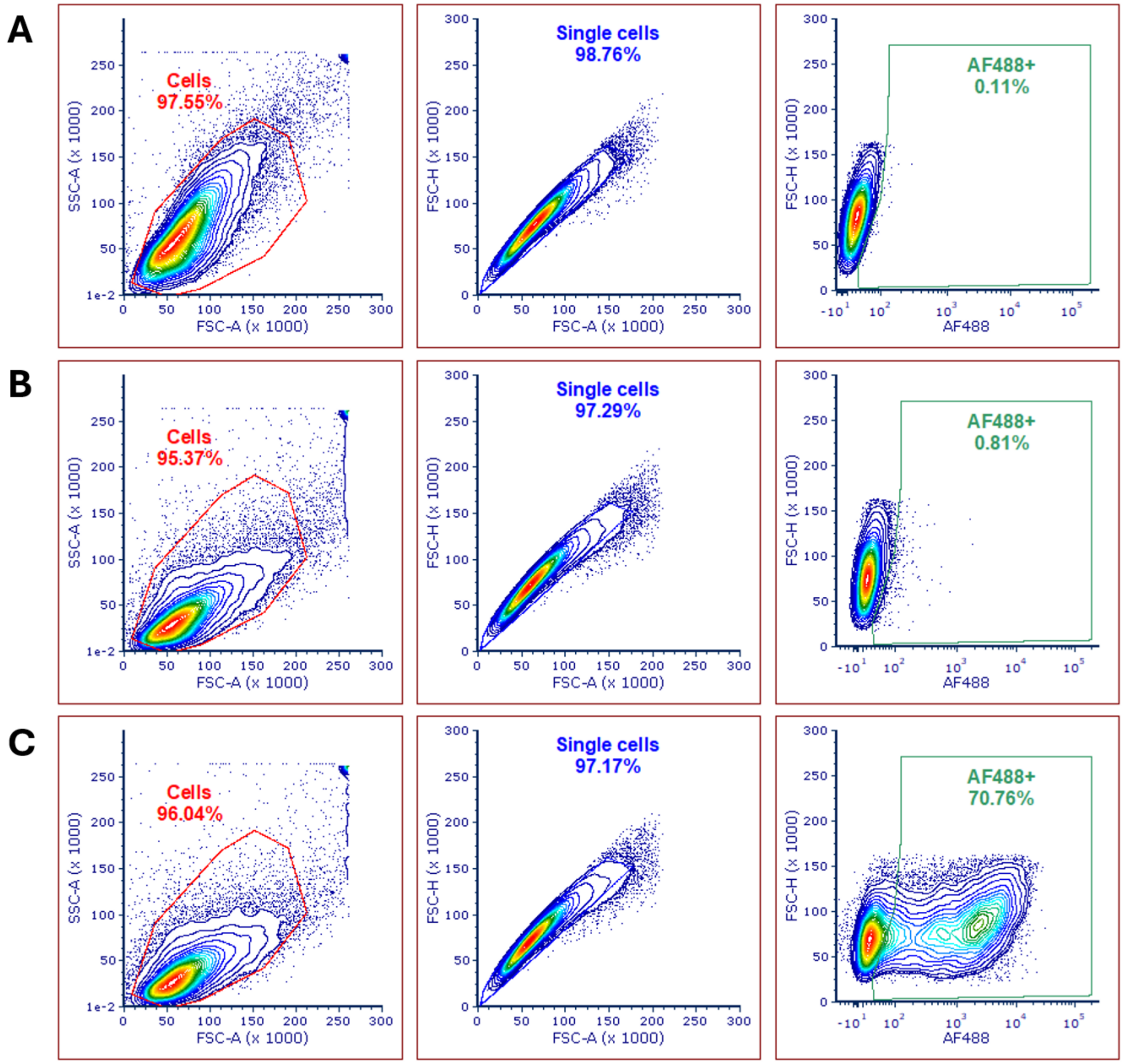


**Fig. S11. Flow cytometry population comparison of AlkNAG labeling in *C. albicans* WT.** DIC185 cells only (A) or DIC185 cells remodeled with 3mM NAG (B) or AlkNAG (C) and then clicked with 20 µM Az488. Flow samples were processed using a BD FACS Aria Fusion Cell Sorter using 50,000 cell counts. Data was processed in FCSExpress 7. The gating method was created using 0.1% TRITC channel signal in the cells only sample and then applied to the experimental samples. Data shown is representative of three biological replicates from at least two technical replicates.

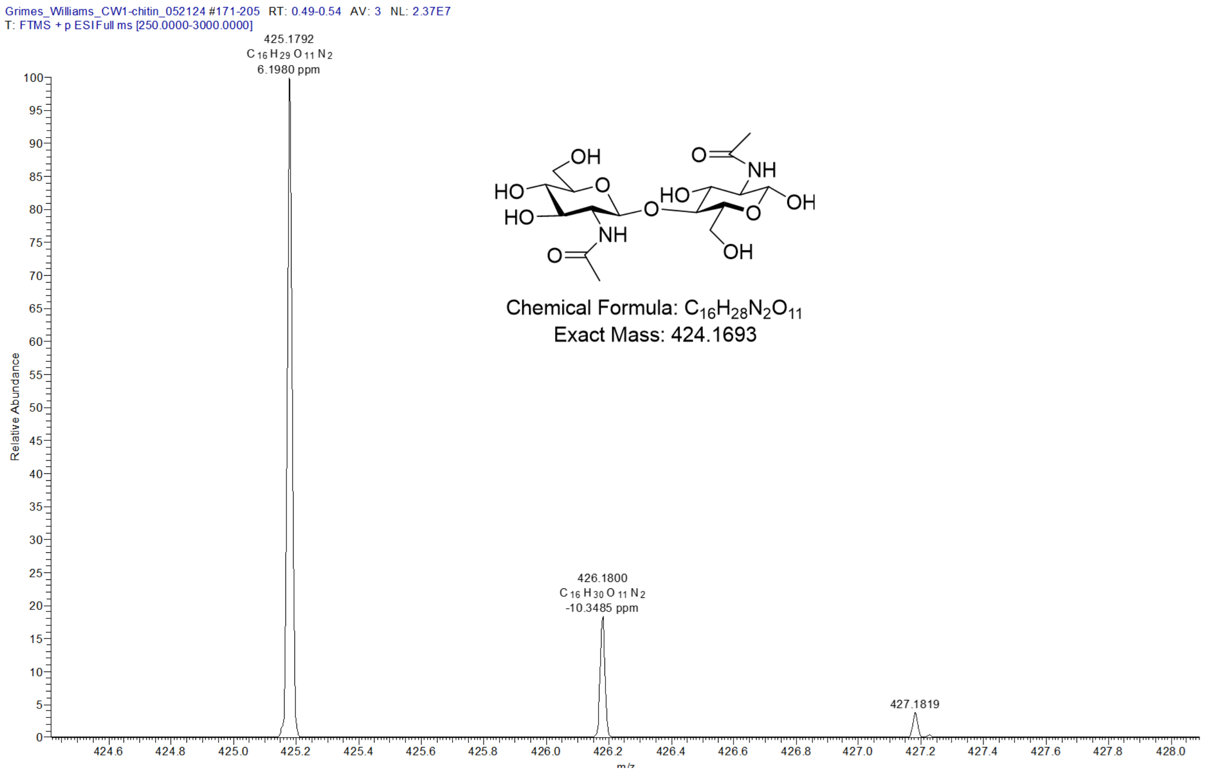


**Fig. S12. HRMS spectra of disaccharide fragment from digested pure chitin from shrimp cells (control).** Chemical structure, expected mass, and observed mass with mass accuracy (ppm) of chitin fragment. The final sample was subjected to high-resolution LCMS, ESI positive mode.

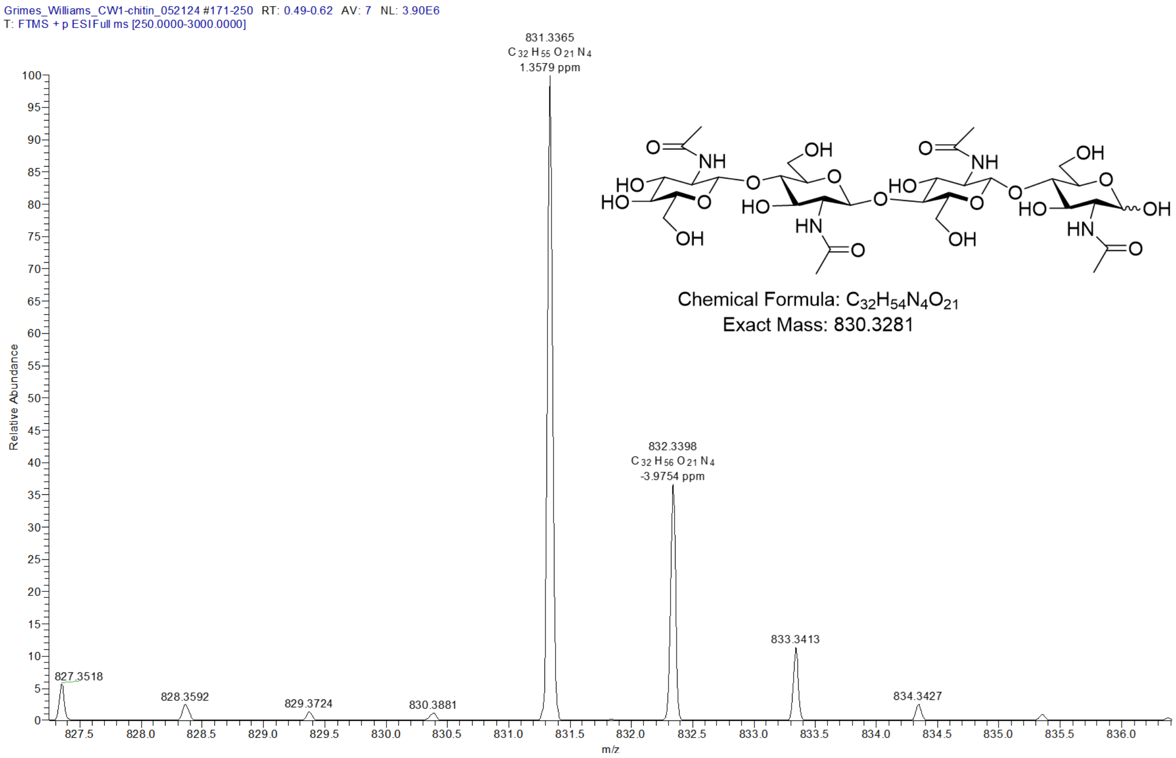


**Fig. S13. HRMS spectra of tetra saccharide fragment from digested pure chitin from shrimp cells (control).** Chemical structure, expected mass, and observed mass with mass accuracy (ppm) of chitin fragment. The final sample was subjected to high-resolution LCMS, ESI positive mode.

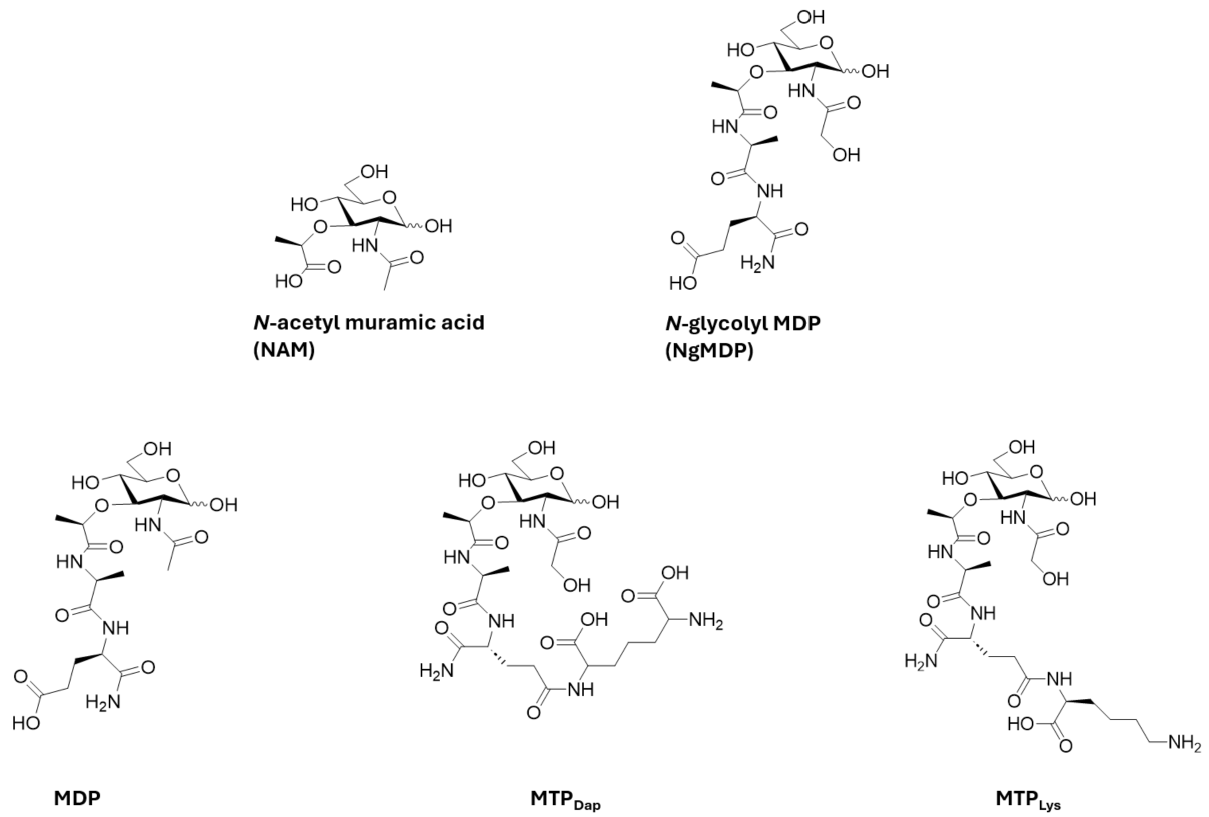


**Fig. S14. Chemical structures of PG synthetic fragments tested in hyphal growth assay.** Compounds MDP, NgMDP, $MTP_{Lys}$, and $MTP_{Dap}$ were purchased commercially from Invivogen. NAM was purchased commercially from Sigma Aldrich.

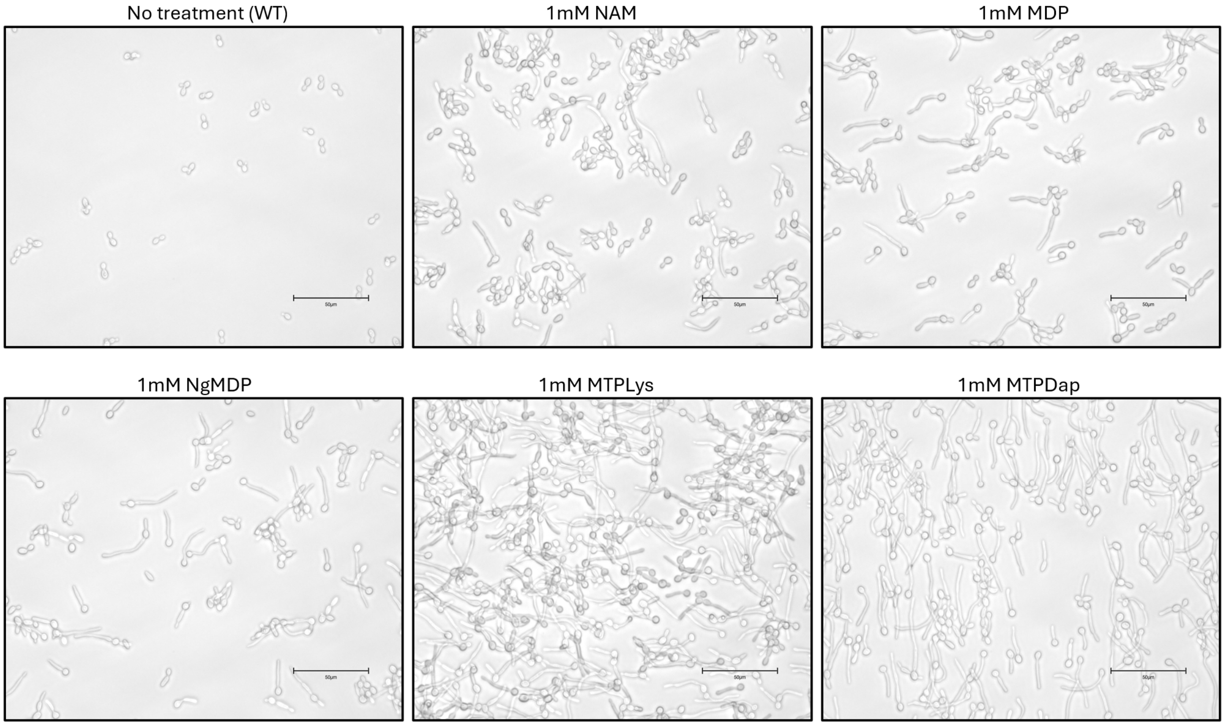


**Fig. S15. Screen of various PG fragment's ability to induce hyphae in *C. albicans* after 4 hours of growth.** Each compound was tested at 1mM final concentration. Scale bars represent 50µM. Experiments were conducted in three biological replicates.

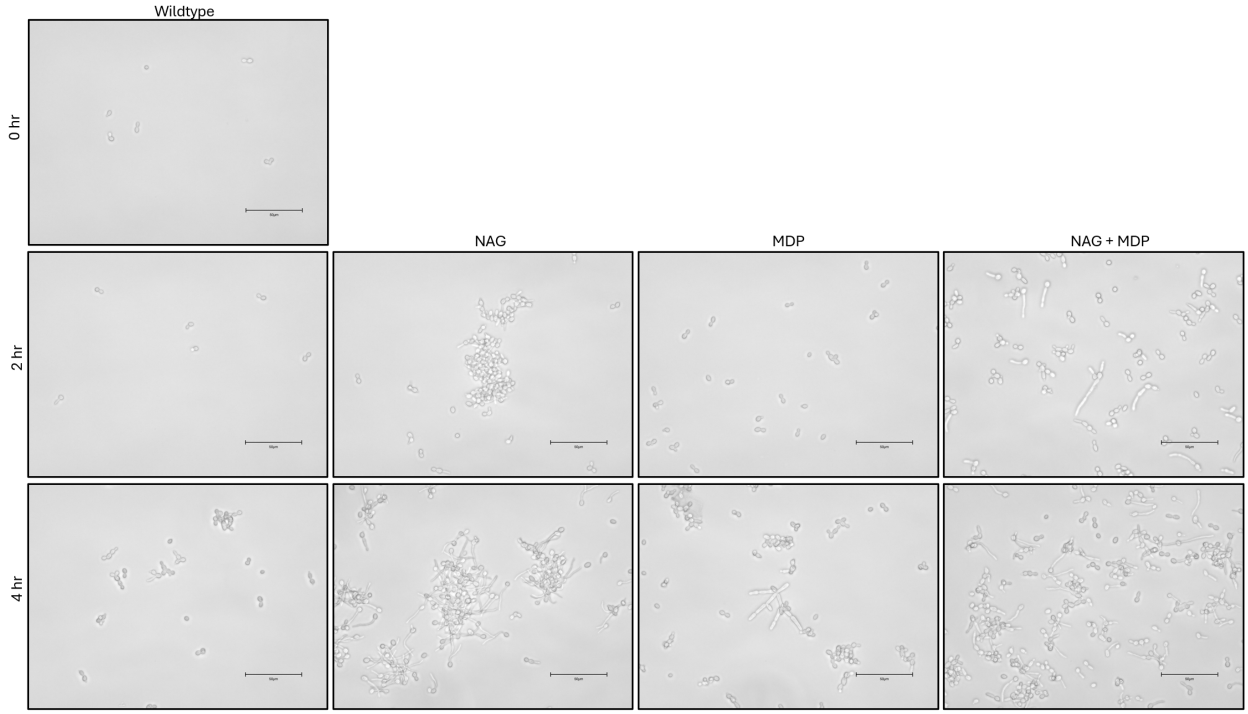


**Fig. S16. Screen of *C. albicans* hyphal growth in response to combinations of various sugars through 4 hours.** Each compound was tested at 1mM final concentration. Scale bars represent 50 µM.

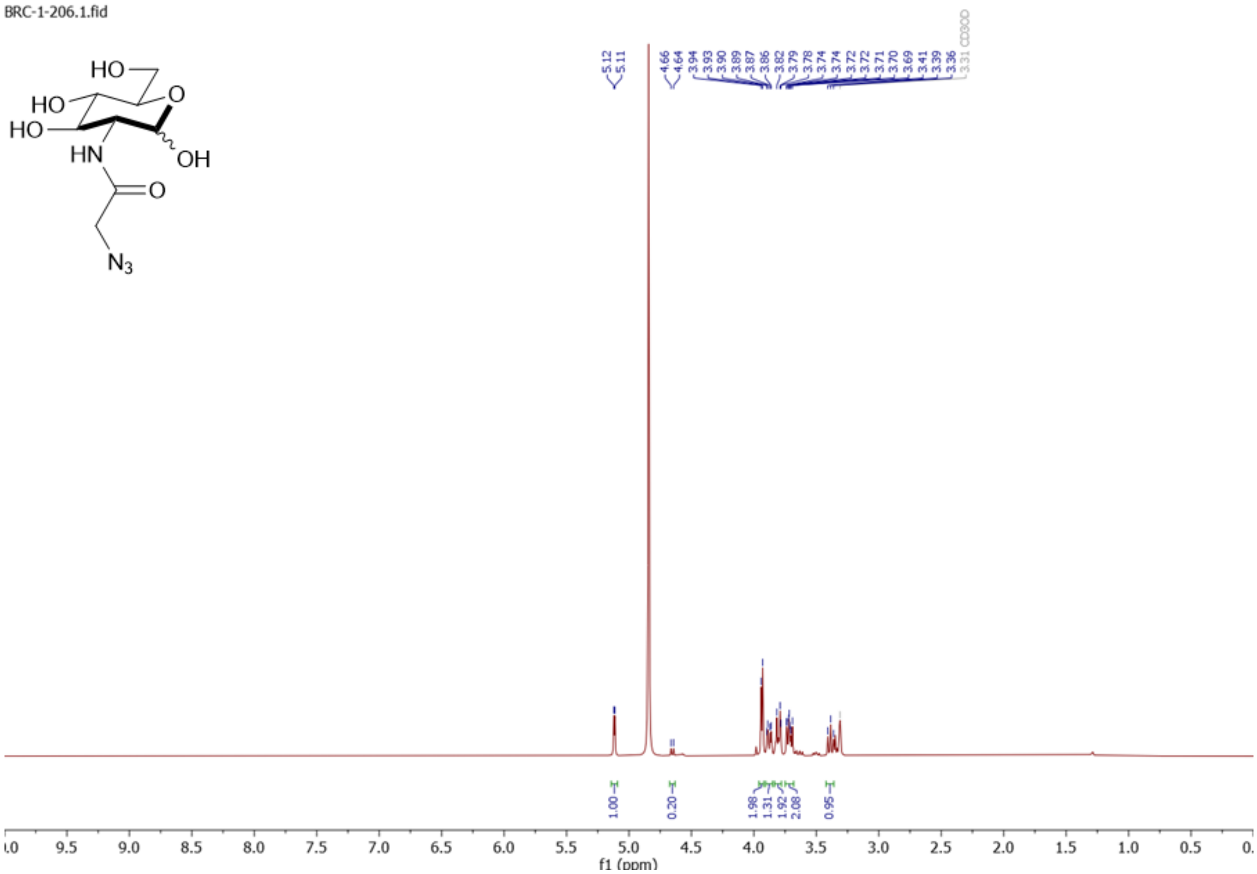


**Fig. S17. $^{1}$H NMR of 2-AzNAG.**

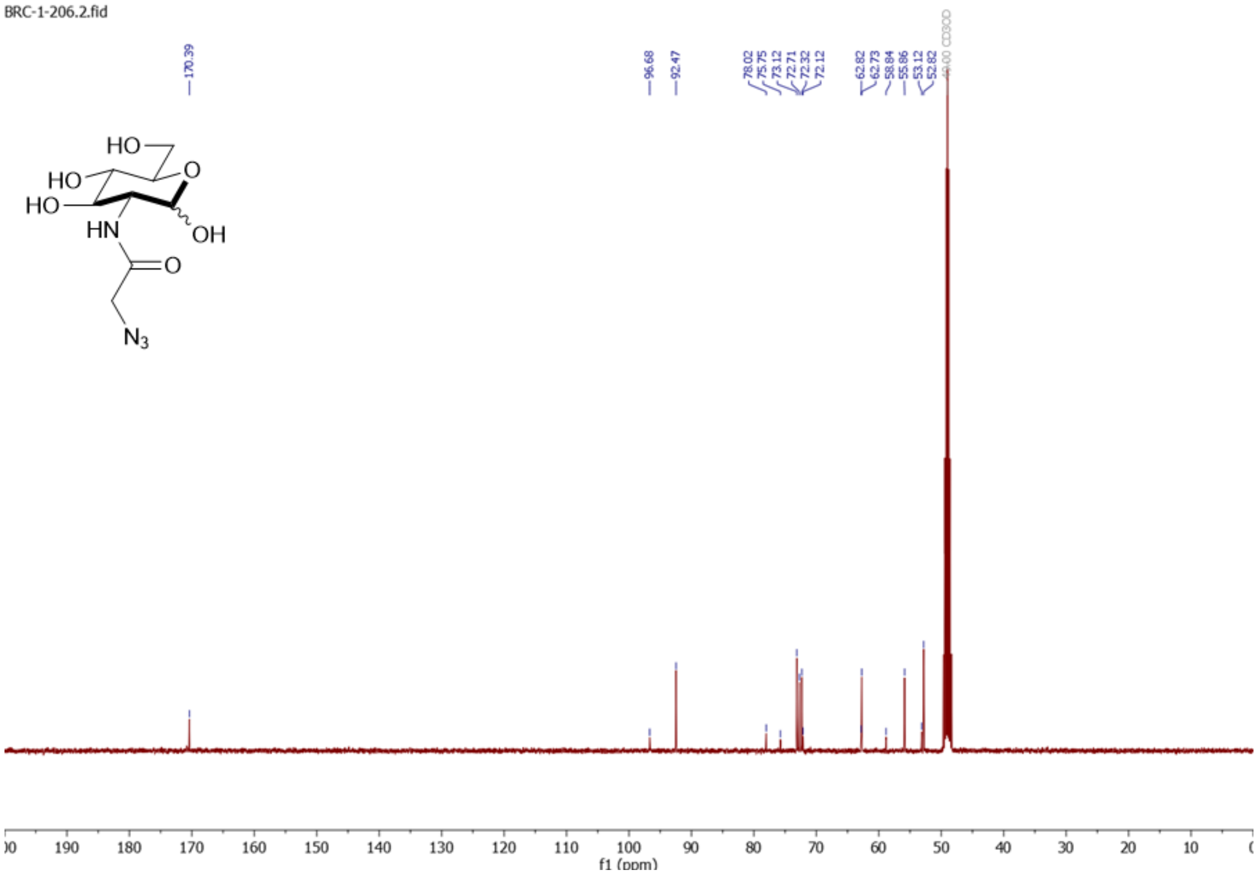


**Fig. S18. $^{13}$C NMR of 2-AzNAG.**

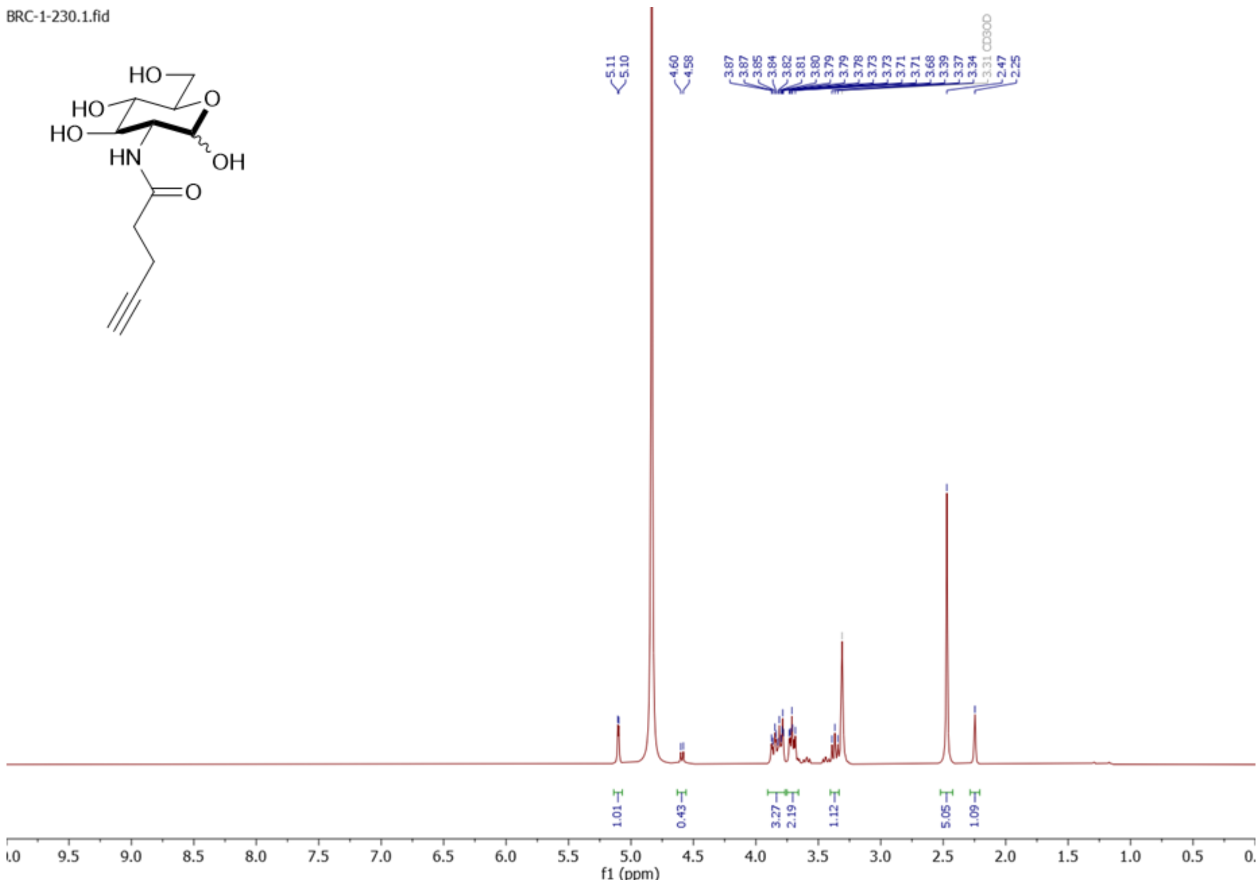

**Fig. S19. $^1$H NMR of 2-AlkNAG.**

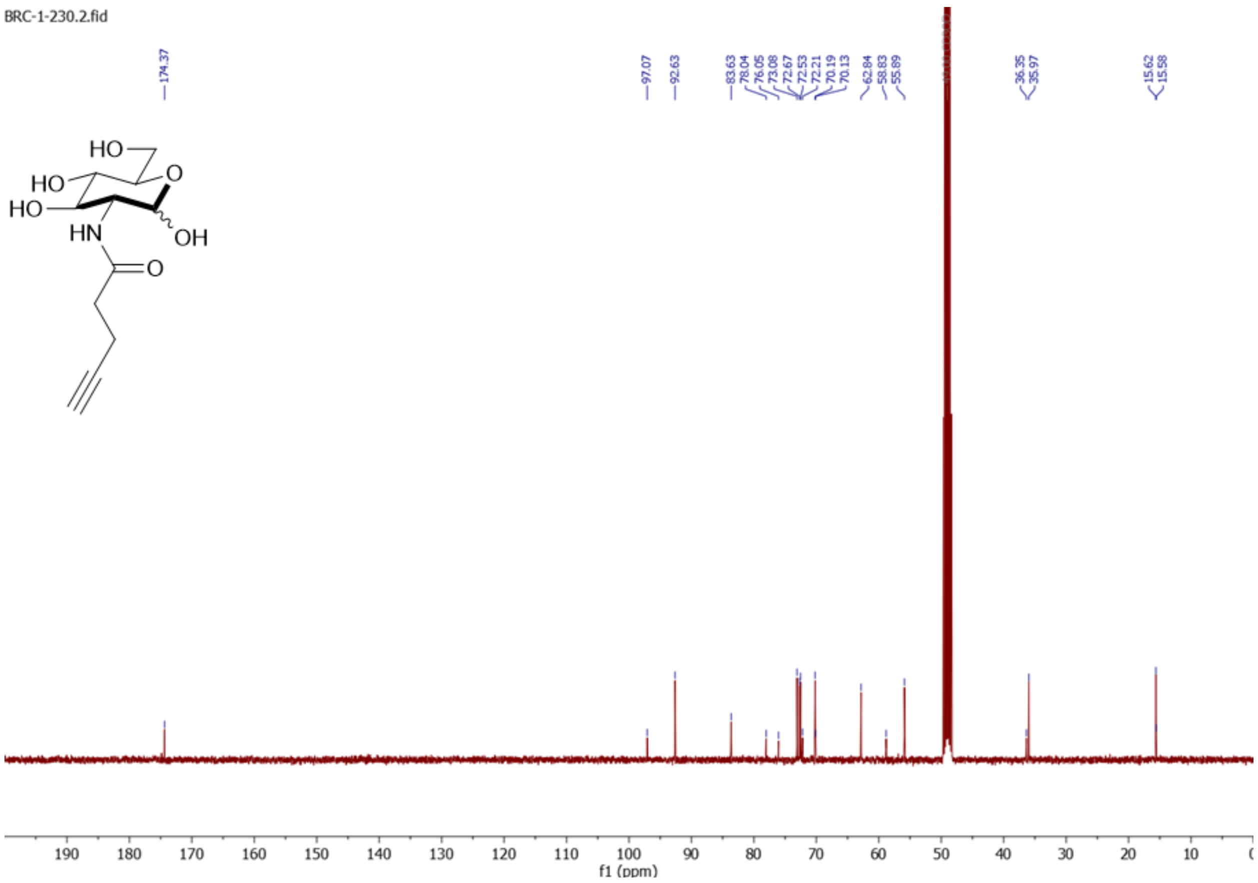


**Fig. S20. $^{13}$C NMR of 2-AlkNAG.**

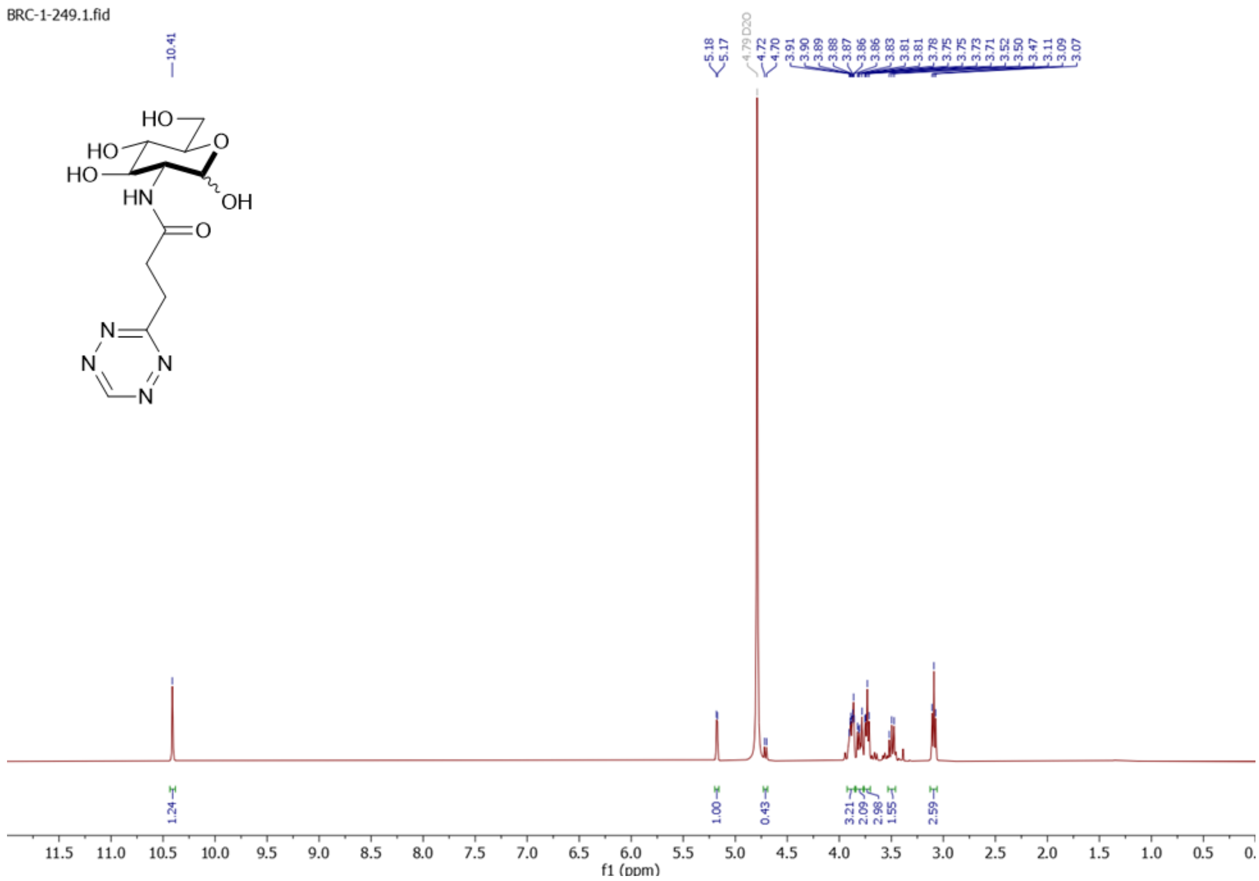


**Fig. S21. $^{1}$H NMR of 2-HTzNAG.**

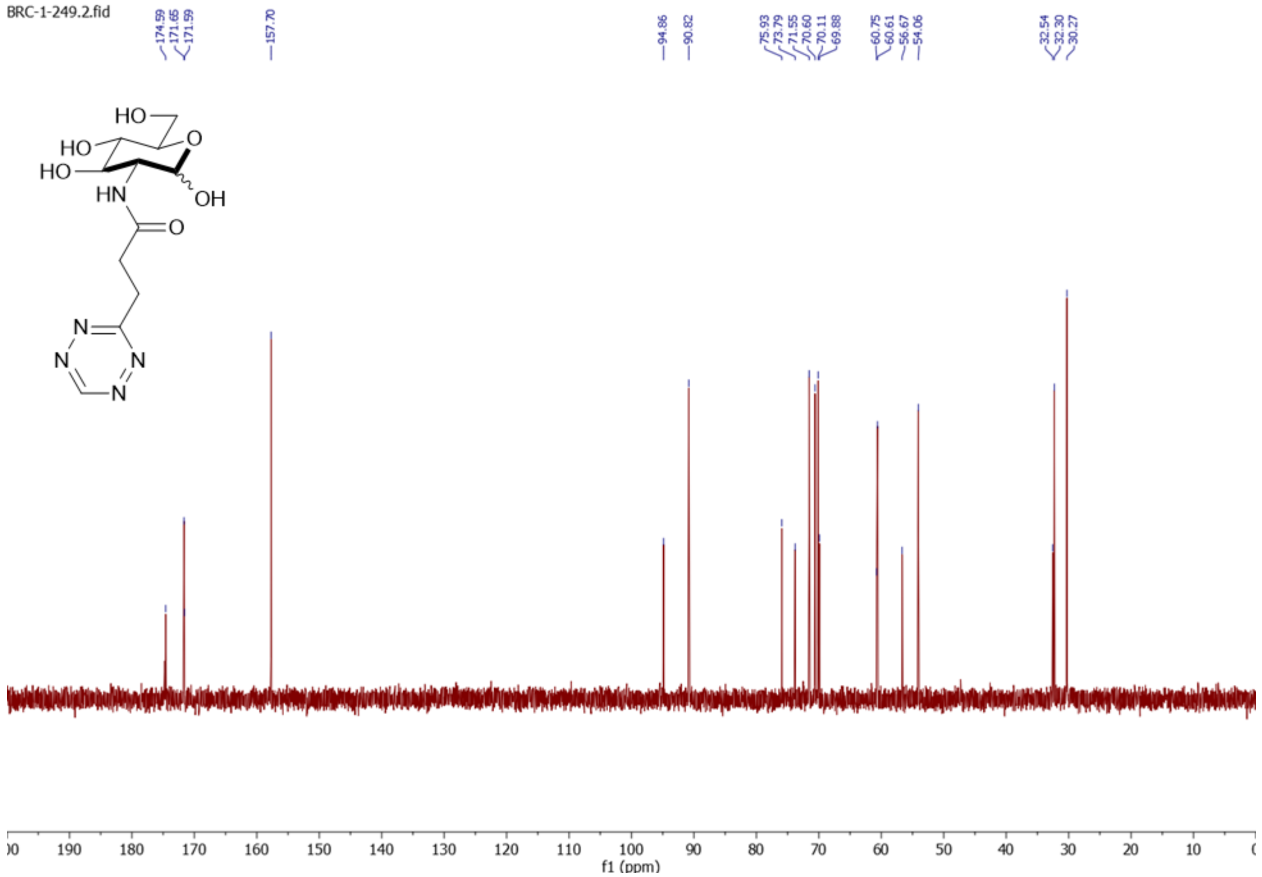


**Fig. S22. $^{13}$C NMR of 2-HTzNAG.**

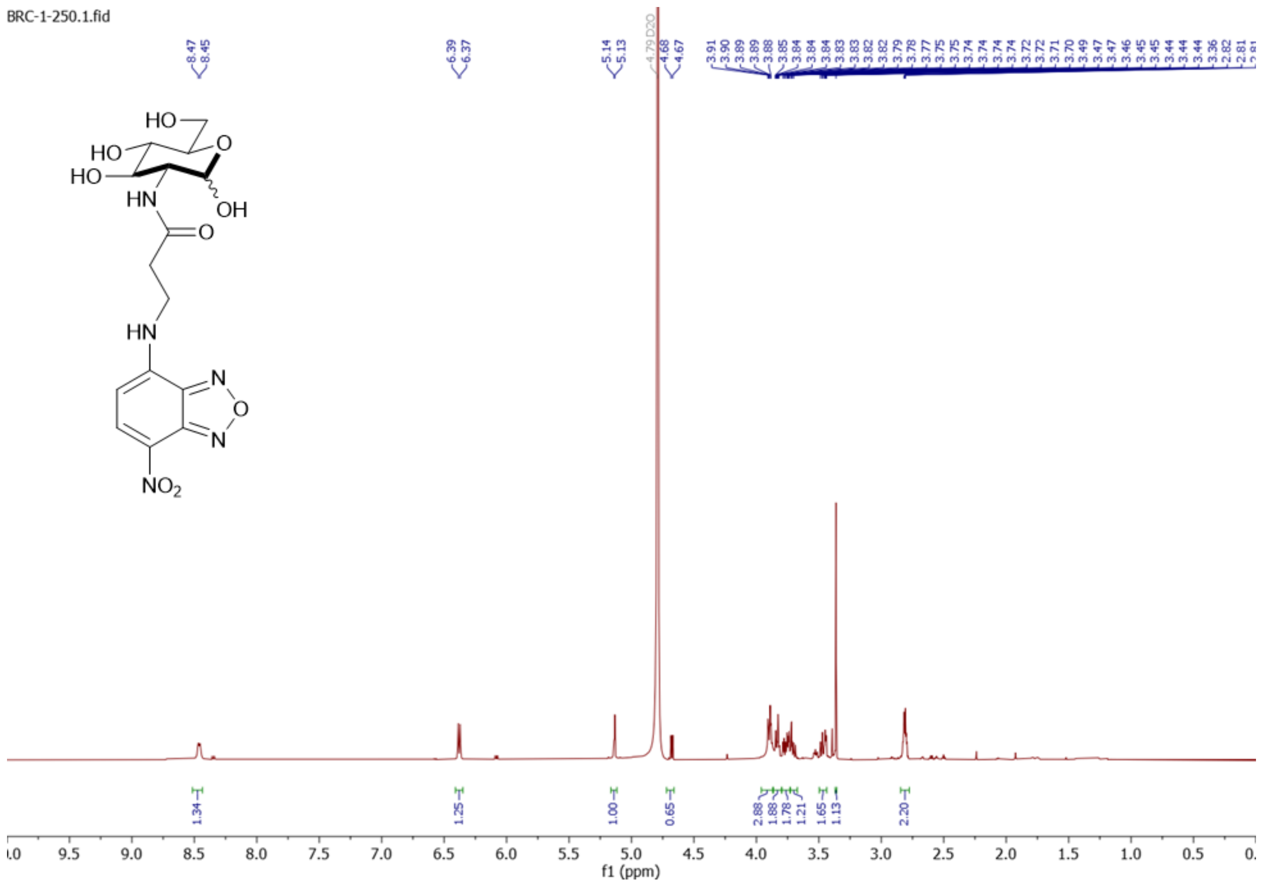

**Fig. S23. [1]H NMR of 2-NBDNAG.**

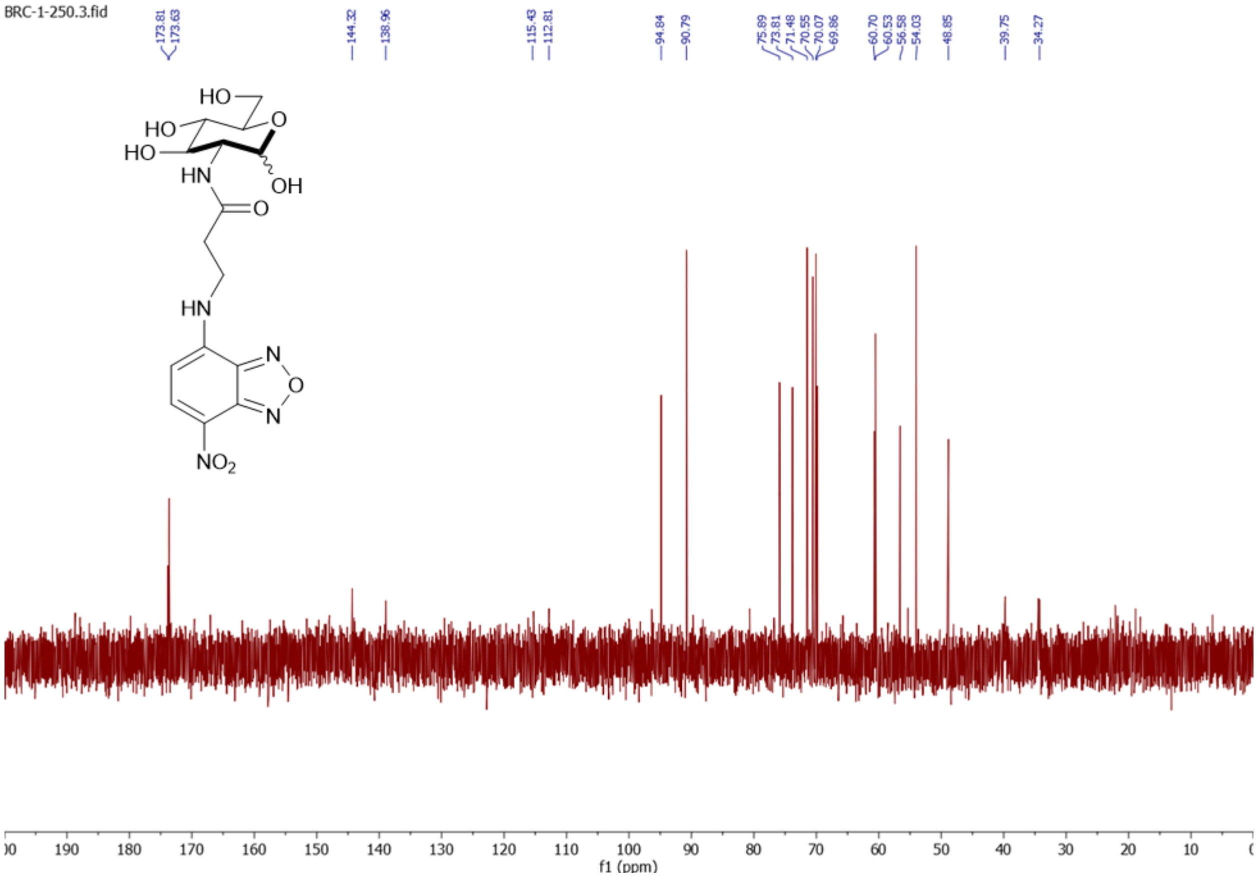


**Fig. S24. $^{13}$C NMR of 2-NBDNAG.**

**Table S1. Characterization of chemoenzymatic analysis of NAG probes.** Detection of products was conducted via High Resolution Mass Spectrometry. Expected mass of product, observed mass of product, and mass accuracy (ppm) of sugars tested.

| Substrate | Theoretical Product m/z $[M-H]^-$ | Experimental Product m/z $[M-H]^-$ | Mass Accuracy (ppm) |
|---|---|---|---|
| GlcNAc | 300.0490 | 300.0481 | -2.9995 |
| Mannose | 259.0224 | 259.0229 | 1.9303 |
| AzNAG | 341.0504 | 341.0506 | 0.5864 |
| AlkNAG | 338.0646 | 338.0652 | 1.7748 |
| HTzNAG | 394.0769 | 394.0774 | 1.2687 |
| NBD-NAG | 492.0773 | 492.0765 | -1.6257 |
| MurNAc | 372.0701 | Not observed | Not observed |
